# Lineage recording reveals hijacked hepatic progenitor states as a common origin of HCC and ICC

**DOI:** 10.64898/2026.07.13.738237

**Authors:** Jinyong Fan, Jiazheng Pei, Nuo Xu, Xinyue Zhang, Xiaoyun Wang, Shiyu Mao, Yu Zhang, Lisi Yu, Yingxia Sun, Yeqing Gong, Xinyuan Xiong, Shu Wang, Lin Li, Zhen Shao, Xuxu Sun, Luonan Chen, Xin Liu

## Abstract

**Highlights:** CERAMIC enables continuous and high-capacity lineage tracing of liver tumor initiation

Fatty liver-associated hepatocytes acquire regenerative and premalignant cell states before malignant transformation

Lineage reconstruction identifies Hep_Bi-zonal cells as the cellular origin of liver tumor initiation

Transcriptional and regulatory programs distinguish tumor-fated hepatocytes from failed-to-transform lineages

Peroxisomal metabolism is required for progenitor-state formation and liver tumor initiation

Spatial remodeling identifies a macrophage niche associated with tumor-fated hepatocytes

Dual ontogenies and functional specialization of lipid-associated macrophages shape the tumor-fated hepatocyte niche

Fatty liver disease predisposes to primary liver cancer, yet the lineage routes and niche mechanisms that select rare tumor-fated hepatocytes remain unclear. Here we developed CERAMIC, a high-capacity CRISPR-Cas9 lineage recorder that co-recovers editing scars and transcriptomes from single cells, and applied it to an AKT/NRAS-driven model of MASLD-associated liver tumor initiation. Longitudinal lineage, single-cell and spatial analyses revealed a hierarchical trajectory in which bipotential bi-zonal hepatocytes (Hep_Bi-zonal), rather than pericentral-like hepatocytes (Hep_CVlike), generated regenerative and neoplastic hepatocyte progenitor states that progressed toward both hepatocellular carcinoma and intrahepatic cholangiocarcinoma lineages. Tumor-fated cells preferentially expanded along a remodeled midlobular-periportal axis and depended on ACOX1-mediated peroxisomal beta-oxidation to withstand lipotoxic and oxidative stress. Spatial and lineage analyses further identified a sequential lipid-associated macrophage niche, in which monocyte-derived LAMs engaged tumor-fated hepatocytes through an LGALS9-P4HB axis, and P4HB inhibition suppressed tumor expansion. These findings define liver tumor initiation as a lineage-restricted process licensed by peroxisomal metabolic adaptation and macrophage-derived niche signals.

## Introduction

Metabolic dysfunction-associated steatotic liver disease (MASLD) is becoming a major driver of primary liver cancer, including hepatocellular carcinoma (HCC) and intrahepatic cholangiocarcinoma (ICC)^1–8^. Its progression to cancer is usually understood as a chronic process involving lipid accumulation, oxidative stress, inflammation, regenerative injury responses and malignant transformation^9–13^. Yet this view leaves a central paradox unresolved. Many hepatocytes experience the same diseased tissue environment, and in experimental models many can be exposed to the same oncogenic pressure, but only a small fraction survives, remodels its identity and enters a tumor-fated trajectory^14–17^. The earliest phase of tumor initiation is therefore not simply determined by when malignant cells first emerge, but by how a small subset of hepatocytes acquires the capacity to persist and undergo clonal expansion before overt tumor formation^18,19^. Understanding this selectivity is essential for defining the cellular logic of fatty liver-associated carcinogenesis.

Recent single-cell and spatial transcriptomic studies have reshaped our understanding of the cellular organization of chronic liver injury and early liver tumorigenesis^20–23^. Hepatocytes are organized along the porto-central lobular axis, where zonated transcriptional and metabolic programs shape homeostatic function, regenerative capacity and injury susceptibility^24–26^. In chronic liver disease and liver cancer, these spatial programs are remodeled, and several studies have identified disease-associated hepatocytes^21^, progenitor-like states^27–30^, onco-fetal neighborhoods^31–34^ and zone-dependent differences in tumorigenic potential^35–39^. These findings suggest that tumor initiation is not simply a consequence of oncogene activation in an otherwise uniform hepatocyte population. They also raise a more precise question: whether tumor-fated cells arise from a pre-existing zonated population^35,38,38,39^, from an injury-induced transitional state^27–29,37,40^, or from stochastic survival among many stressed hepatocytes^41^. However, transcriptomic state maps alone cannot determine whether a given hepatocyte state is a true ancestor of malignant cells, a transient repair state, a stress-induced dead end or a parallel response to tissue damage^21,42–44^. The field therefore needs lineage-resolved evidence that links hepatocyte state to future tumor fate^45–48^.

Lineage tracing provides a direct route to obtaining such evidence^45^. CRISPR-based lineage-recording systems have enabled reconstruction of cellular histories during development, regeneration and cancer progression, particularly when lineage scars can be recovered together with single-cell transcriptomes^49–58^. Tumor initiation poses particularly stringent requirements for lineage recording because the cells that ultimately acquire malignant fate are both rare and transient. A suitable recorder must continuously capture these transitional cells over time, generate sufficient barcode diversity to resolve expanding clones, and preserve lineage information while simultaneously enabling accurate definition of cell states across complex tissue environments. ^45,59^. Existing recorders have established the power of this strategy but can be constrained by limited target-site number, intersite deletions that erase prior information, or inefficient recovery in standard single-cell workflows^49,56,60^. These constraints are especially problematic when the biological question is not only which clone expands, but which early cell state gives rise to that clone. A recorder suited to this problem must therefore connect three layers in the same cell: accumulated lineage history, current transcriptional identity and the tissue context in which fate decisions occur^57,61^.

Cell-intrinsic lineage fate is only one part of the problem. Fatty liver-associated tumor initiation occurs within a metabolically stressed and immunologically remodeled tissue, where macrophages are prominent responders to lipid accumulation, cell death and repair^62–65^. Lipid-associated macrophages (LAMs)^65,66^ have been implicated in metabolic disease^43,67^, fibrosis regression^68,69^, tissue repair^67,70–72^ and tumor ecosystems^73,74^, but their functions are context-dependent. In liver injury, both resident Kupffer cells and recruited monocytes can contribute to macrophage remodeling, raising the possibility that macrophage ontogeny and timing shape distinct phases of the tumor-promoting niche^75–77^. This distinction is important because macrophages that share lipid-handling markers may differ in origin, spatial localization and functional output^78–80^. Whether such macrophage populations merely accompany damaged hepatocytes or provide signals that support tumor-fated hepatocyte expansion during initiation remains unresolved.

Here we developed CERAMIC, a CRISPR-Cas9 Evolving Recording Arrays Multiplexed by Csy4 platform designed to generate high-complexity lineage barcodes that are co-captured with endogenous transcriptomes. We applied CERAMIC to an AKT/NRAS-driven model of MASLD-associated liver tumor initiation^81,82^ and integrated longitudinal single-cell transcriptomics, spatial transcriptomics, metabolic perturbation and macrophage lineage tracing. This strategy allowed us to reconstruct the lineage hierarchy of tumor-fated hepatocytes, define the metabolic program that permits their survival and progenitor-state transition, and map the macrophage niche that supports their expansion. Our results identify Hep_Bi-zonal cells as a major tumor-fated hepatocyte population and reveal that ACOX1-dependent peroxisomal adaptation and monocyte-derived LAM-associated signaling cooperate to license early liver tumor initiation.

## Results

### CERAMIC enables continuous and high-capacity lineage tracing of liver tumor initiation

To reconstruct lineage history and cell-state dynamics during liver tumor initiation, we developed CERAMIC (CRISPR-Cas9 Evolving Recording Arrays Multiplexed by Csy4), a high-capacity lineage-recording platform that enables simultaneous recovery of lineage information and transcriptomic state from single-cell RNA sequencing (Figure 1a). CERAMIC was rationally engineered to maximize barcode complexity while minimizing lineage information loss during long-term recording. Specifically, the recorder incorporates 24 independently expressed sgRNAs together with a synthetic target-site array organized into eight recording modules (TA1-TA8) spanning more than 3 kb, which enable long-term recording of hepatocyte fate transitions (Figure 1a; Supplementary Table 1). Each target site was uniquely matched to one sgRNA, enabling parallel accumulation of editing scars across the array (Figure 1a). Because large intersite deletions represent a major source of lineage information loss in existing CRISPR recorders^56,60,83^, we substantially increased the spacing between adjacent target sites to approximately 40 bp, nearly tenfold longer than in previous designs. In addition, every three target sites were strategically followed by a 30-nt poly(A) sequence and a 28-nt Csy4 recognition sequence^84,85^, thereby partitioning the recorder into discrete recording modules (Figure 1a). This modular architecture enables each recording block to function as an independent recording unit while substantially reducing the probability that large intersite deletions erase previously accumulated lineage information. (Figure 1b, c). Following transcription, the full-length target array is processed by Csy4 into ∼300 bp barcode-containing transcripts^86^, enabling co-capture of lineage information and endogenous gene-expression profiles within single-cell RNA sequencing workflows (Figure 1a). To enable inducible lineage recoding, Cas9 expression was placed under doxycycline control, whereas the recorder array, Csy4, and TetOn-3G components were constitutively expressed from EF1a-driven cassettes (Figure 1a). We first established the recorder configuration in embryonic stem cells and subsequently generated CERAMIC mice by integrating the sgRNA cassette into the H11 safe-harbor locus and the remaining recorder components into the Rosa26 locus. Crossing these strains produced animals carrying all transgenic elements required for inducible lineage recording (Supplementary Table 1).

**Figure 1.**
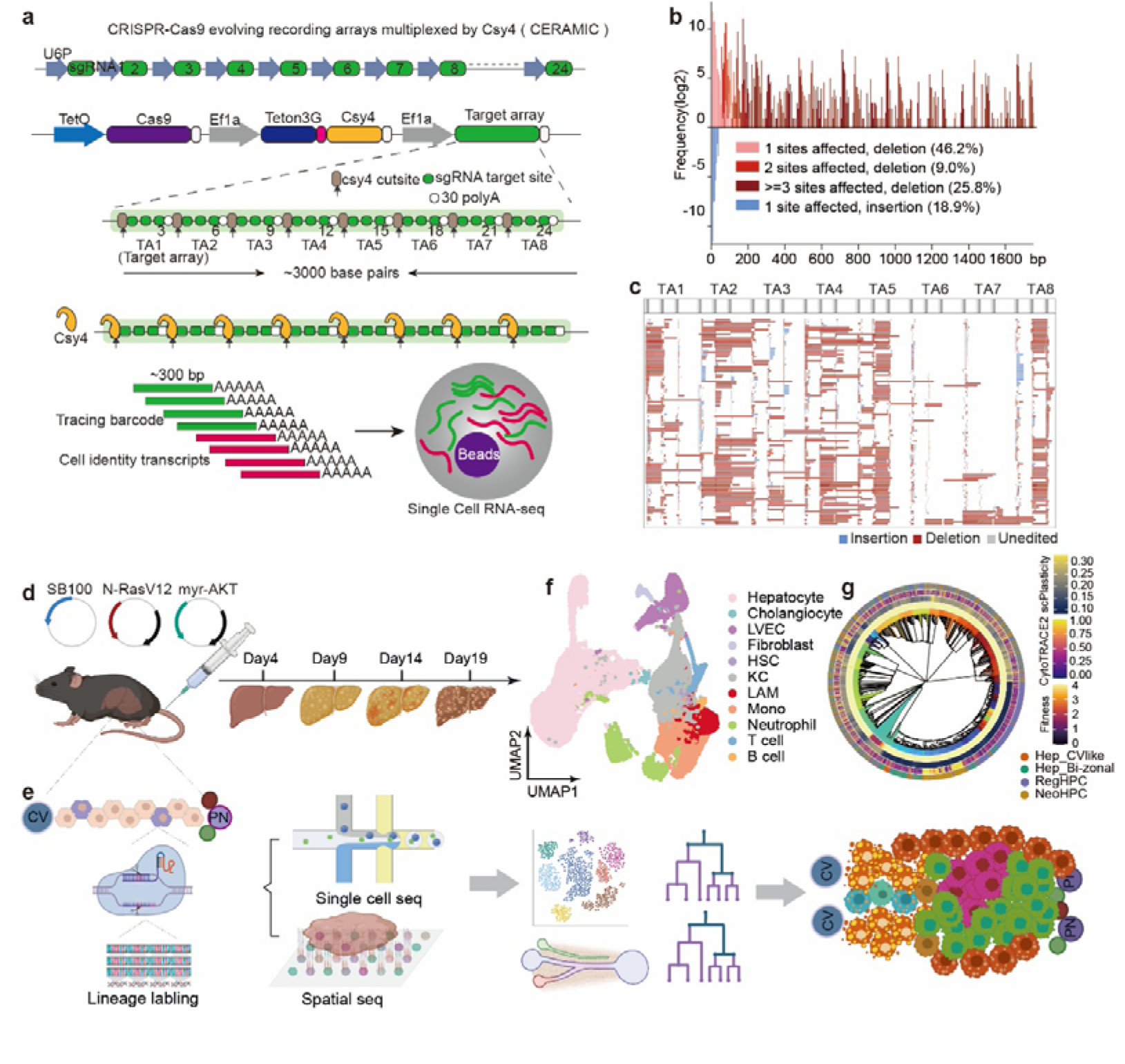
Design and in vivo application of the CERAMIC lineage-recording system for tracking liver tumor initiation. **(a)** Schematic design of the CRISPR-Cas9 Evolving Recording Arrays Multiplexed by Csy4 (CERAMIC) lineage-recording system. A multiplexed array of 24 sgRNAs is driven by a U6P promoter. The core recording construct features a doxycycline-inducible Cas9 cassette (under TetO control), an Ef1a-driven Teton3G transactivator, a Csy4 endoribonuclease, and a ∼3000-bp synthetic target array comprising eight tandem modules (TA1-TA8). Each TA module is engineered with a Csy4 cleavage site, an sgRNA target site, and a poly(A) capture element. Subsequent Csy4 processing cleaves the continuous array transcript into individual ∼300-bp barcode transcripts, enabling co-capture of lineage records alongside endogenous cellular mRNAs via droplet-based single-cell RNA sequencing (scRNA-seq). **(b)** Frequency distribution of editing outcomes recovered from CERAMIC mouse embryonic stem cells (ESCs) after 48 h of induction with 100 ng/mL Dox, classified by the number of affected target sites and mutation types. The y-axis represents the log2-transformed frequency of each editing event, while the x-axis indicates the size of insertions or deletions (in base pairs, bp). Color-coded bars distinguish single- and multi-target-site deletions (above the baseline) and single-target-site insertions (below the baseline). **(c)** Representative editing profiles across the target-site modules (TA1-TA8). A consolidated profile of 100 representative edited CERAMIC alleles generated in CERAMIC mouse embryonic stem cells (ESCs) after 48 h of induction with 100 ng/mL doxycycline (Dox), compiled from the experiments shown in Supplementary Fig. S1a and S1e. Each row represents an individual edited allele, with insertions, deletions, and unedited regions shown in blue, red, and light gray, respectively. In the schematic profile positioned above the allele matrix, dark gray bars demarcate the precise double-strand break (DSB) windows targeted by the sgRNAs, while the light gray regions define the broader windows permissible for genetic editing. **(d)** Experimental workflow for lineage-resolved profiling of liver tumor initiation. CERAMIC mice are subjected to hydrodynamic tail-vein injection of three distinct transposon plasmids: the SB100 plasmid (blue vector) expressing the SB100 transposase protein, the N-RasV12 oncogene co-expressed with a red fluorescent protein on a single vector (red vector), and the myr-AKT oncogene co-expressed with a green fluorescent protein on a single vector (green vector). Black segments within the vectors represent the oncogene sequences. Liver tissues were collected at Days 4, 9, 14, and 19 after hydrodynamic tail-vein injection. **(e)** Integrated multi-omics strategy to track tumor evolution spanning the hepatic lobule architecture from the central vein (CV) to the portal node (PN). Following hydrodynamic tail-vein injection of the oncogenes into CERAMIC mice, a subset of hepatocytes along the CV-PN axis integrates the oncogenes (represented as purple hepatocytes), while other hepatocytes remain unintegrated (represented as pink hepatocytes). Upon subsequent doxycycline induction, Cas9 expression is activated within the cells, forming a complex with the sgRNAs to cleave the target arrays and generate heritable genetic scars. Double-positive oncogene-expressing hepatocytes are sequentially harvested at successive experimental time points for single-cell RNA sequencing, while parallel liver tissue sections are processed for spatial transcriptomics. Single-cell RNA sequencing and spatial transcriptomics were performed at the indicated time points, and the resulting datasets were used for downstream lineage and spatial analyses. PN = portal node (a vasculature structure composed from the portal vein (purple), hepatic artery (red) and bile duct (green)), CV = central vein (blue). **(f)** UMAP embedding of the integrated single-cell cellular atlas, resolving major hepatic, stromal, and immune populations within the tumor microenvironment, including hepatocytes, cholangiocytes, liver vascular endothelial cells (LVECs), fibroblasts, hepatic stellate cells (HSCs), Kupffer cells (KCs), lipid-associated macrophages (LAMs), monocytes, neutrophils, T cells, and B cells. **(g)** Representative reconstructed lineage phylogeny and spatial clonal architecture at Day 4 post-induction integrated with functional single-cell parameters. The circular phylogenetic tree is organized into multi-layered concentric rings mapped to individual cells: the innermost layer displays a rainbow of distinct colors representing individual clones; the second layer indicates the inferred clonal growth advantage (Fitness score); the third layer reflects the single-cell developmental potential (scPlasticity); the fourth layer shows the CytoTRACE2 differentiation score; and the outermost layer specifies the corresponding cell types across the tree.

Next, we evaluated the ability of CERAMIC to generate inducible and highly diverse lineage barcodes in embryonic stem cells following doxycycline (Dox) induction. Bulk genomic DNA was collected at the indicated time points, and barcode arrays were amplified and subjected to targeted amplicon sequencing, followed by reference alignment and allele calling as described in Methods (Supplementary Table 2). Robust, dose-dependent accumulation of editing scars was observed across all target arrays, with editing efficiencies ranging from approximately 50% to 90%, depending on the target site. (Supplementary Figure 1a; Supplementary Table 2). Across the five biological replicates, the vast majority of editing scars were replicate-specific, with individual experiments containing 5,432, 5,232, 4,116, 3,250, and 1,253 unique barcode species, respectively. Across biological replicates, fewer than 300 barcode species were shared between any pair of samples, whereas the vast majority of barcode species were unique to individual replicates (Supplementary Figure 1b). Together, these results demonstrate that CERAMIC generates highly diverse and largely non-overlapping barcode repertoires across independent experiments, providing sufficient barcode complexity for high-resolution lineage reconstruction.

Editing spectrum revealed a broad range of repair outcomes, including deletions spanning 1-1,750 bp and insertions of up to 21 bp in length (Figure 1b, c; Supplementary Table 2). Among all editing events, 55.2% were confined to deletions spanning one or two adjacent sgRNA target sites (46.2% affecting a single site and 9.0% affecting two sites), while insertion events accounted for 18.9% of the total repair outcomes. By contrast, large deletions spanning three or more sgRNA target sites accounted for 25.8% of all events (Figure 1b, c), demonstrating that modular partitioning effectively constrains long-range deletions and preserves accumulated lineage information during continuous recording.

We further quantified the diversity of editing scars generated at individual target sites. Across the 24 recording sites, each sgRNA produced approximately 22,031-63,130 distinct editing patterns, with an average of 34,344 unique edit modes per site, demonstrating extensive mutational diversity throughout the recorder array (Supplementary Figure 1c; Supplementary Table 2). In total, we observed 824,252 site-resolved editing pattern records, corresponding to 698,872 globally unique editing patterns after merging identical patterns across sites. These editing scars were dominated by intrasite indels (26.5-48.8%) and intrasite deletions (25.5-38.0%), while insertion-only patterns accounted for 4.0-14.3% of patterns across sites. Intersite deletions represented 3.7-33.3% of unique editing patterns, whereas intersite indels were rare, accounting for only 0.01-0.26% (Supplementary Figure 1d). Together, these results demonstrate that CERAMIC generates highly diverse and information-rich lineage barcodes with substantial combinatorial complexity, enabling scalable and long-term phylogenetic reconstruction.

A key requirement for lineage-recording systems is the ability to continuously generate and accumulate heritable information throughout cell proliferation, thereby enabling long-term tracking of lineage relationships rather than producing static lineage labels^46,54,56,87^. To directly assess the long-term recording capacity of CERAMIC, we established a serial-passaging assay using recorder-engineered embryonic stem cells and continuously monitored editing dynamics over nine consecutive passages spanning 12 days (Supplementary Figure 1e; Supplementary Table 2). Longitudinal sequencing revealed the sustained emergence of newly generated editing scars throughout the entire recording period, indicating persistent recorder activity across multiple rounds of cell division (Supplementary Figure 1f, g). Importantly, new scars accumulated progressively while existing scars were preserved, allowing lineage information to accumulate cumulatively across successive passages. Recording activity was sustained throughout the experimental period without detectable saturation as evidenced by the continued appearance of newly generated editing scars through the final passage, suggesting that CERAMIC is capable of continuously labeling proliferating cells over extended timescales. Together, these findings establish CERAMIC as a durable lineage recorder capable of continuously accumulating heritable lineage information over extended periods of cell proliferation.

Following transcription, the full-length target-site array is processed by Csy4 into ∼300-bp poly(A)-containing barcode transcripts, enabling efficient co-capture of lineage barcodes together with endogenous mRNAs by standard droplet-based single-cell RNA sequencing and thereby simultaneous recovery of lineage history and cellular state from the same cell (Figure 1a; see Methods). We next evaluated the single-cell recording performance of CERAMIC using an in vitro reprogramming model established from fibroblasts isolated from E13.5 CERAMIC embryos. Across three independent single-cell RNA-sequencing experiments, lineage barcodes were successfully recovered from more than 90% of high-quality cells, demonstrating highly efficient co-capture of lineage information together with endogenous transcriptomes (Supplementary Figure 1h). To further assess recorder completeness, we examined recovery of the eight independent recording modules, each comprising three sgRNA target sites. Overall, 90.2% of barcode-positive cells retained all eight recording modules, whereas only 8.3% contained partially recovered arrays, indicating that CERAMIC enables near-complete recorder recovery at single-cell resolution (Supplementary Figure 1i). Collectively, these results demonstrate that CERAMIC supports robust single-cell lineage recording with both highly efficient barcode recovery and faithful preservation of recorder integrity, thereby providing sufficient lineage information for accurate phylogenetic reconstruction while simultaneously resolving cellular states.

We then assessed recorder performance in vivo. Doxycycline administration induced robust Cas9 expression across multiple organs, resulting in efficient editing of the target array with tissue-specific editing efficiencies ranging from 60%-80% (Supplementary Figure 1j, k; see Methods). In contrast, uninduced animals displayed minimal background editing (∼1%), indicating tight temporal control of recorder activity. Together, these results establish CERAMIC as a versatile platform capable of generating highly diverse and continuously evolving lineage barcodes in vivo.

Having established that CERAMIC enables efficient and faithful reconstruction of lineage histories at single-cell resolution, we next applied this platform to investigate how hepatocytes undergo fate transitions during MASLD-associated liver tumor initiation. We combined CERAMIC lineage recording with an AKT-NRAS-driven mouse model generated by hydrodynamic tail-vein injection (HTVI), which rapidly induces hepatic steatosis followed by the development of mixed hepatocellular carcinoma (HCC) and intrahepatic cholangiocarcinoma (ICC) within approximately one month ^81,82^(Figure 1d; see Methods).

To comprehensively capture the cellular and lineage dynamics throughout tumor initiation, we performed longitudinal single-cell RNA sequencing and spatial transcriptomic profiling of liver tissues collected at baseline and successive stages following AKT-NRAS induction (Days 4, 7, 9, 14, and 19) (Figure 1d, e). This integrated atlas resolved all major hepatic, stromal, and immune cell populations, including hepatocytes, cholangiocytes, liver vascular endothelial cells (LVECs), fibroblasts, hepatic stellate cells (HSCs), Kupffer cells (KCs), lipid-associated macrophages (LAMs), monocytes, neutrophils, T cells, and B cells (Figure 1f; Supplementary Figure 2a, b; Supplementary Table 3). Mapping of AKT and NRAS transgene expression further distinguished oncogene-expressing hepatocyte-derived populations from non-transformed hepatocytes, thereby establishing a lineage-resolved framework for tracking hepatocyte fate transitions throughout tumor initiation (Figure 1f; Supplementary Figure 2c, d).

CERAMIC lineage information was broadly recovered across the entire in vivo cellular atlas. Among 80,530 transcriptome-qualified cells, 56,906 cells contained detectable lineage barcodes, corresponding to an overall barcode recovery rate of 70.7% (Supplementary Figure 2g, h; Supplementary Table 3). Of these barcode-positive cells, 43,388 contained CRISPR-generated editing scars, representing 53.9% of all transcriptome-qualified cells and 76.2% of barcode-positive cells (Supplementary Figure 2f, h). the majority of recovered barcodes contained informative editing scars suitable for downstream phylogenetic reconstruction. Notably, barcode recovery was broadly observed across hepatic, stromal, and immune compartments, indicating that lineage information was preserved throughout diverse cellular states rather than being restricted to specific cell populations. Barcode-positive cells retained an average of 2.53 recovered recording modules, corresponding to more than six sgRNA target sites per cell, with a median of two recovered target-array units (Supplementary Figure 2e, h). Together, these findings establish that CERAMIC provides robust in vivo lineage recording with sufficient barcode complexity and recorder recovery to enable reliable phylogenetic reconstruction during liver tumor initiation.

### Fatty liver-associated hepatocytes acquire regenerative and premalignant cell states before malignant transformation

To define hepatocyte states associated with liver tumor initiation, we extracted normal hepatocytes together with oncogene-expressing hepatocyte-derived cells from the integrated atlas and performed high-resolution re-clustering (Figure 2a; Supplementary Tables 3 and 4). This analysis resolved seven transcriptionally distinct hepatocyte states based on integrated clustering together with day-specific UMAP visualization (see Methods), including normal hepatocytes (Hep_Normal), central vein-like hepatocytes (Hep_CVlike), and bi-zonal hepatocytes (Hep_Bi-zonal), regenerative hepatocyte progenitor cells (RegHPC), neoplastic hepatocyte progenitor cells (NeoHPC), hepatocellular carcinoma (HCC), and intrahepatic cholangiocarcinoma (ICC) (Figure 2a; Supplementary Figure 2c, d).

**Figure 2.**
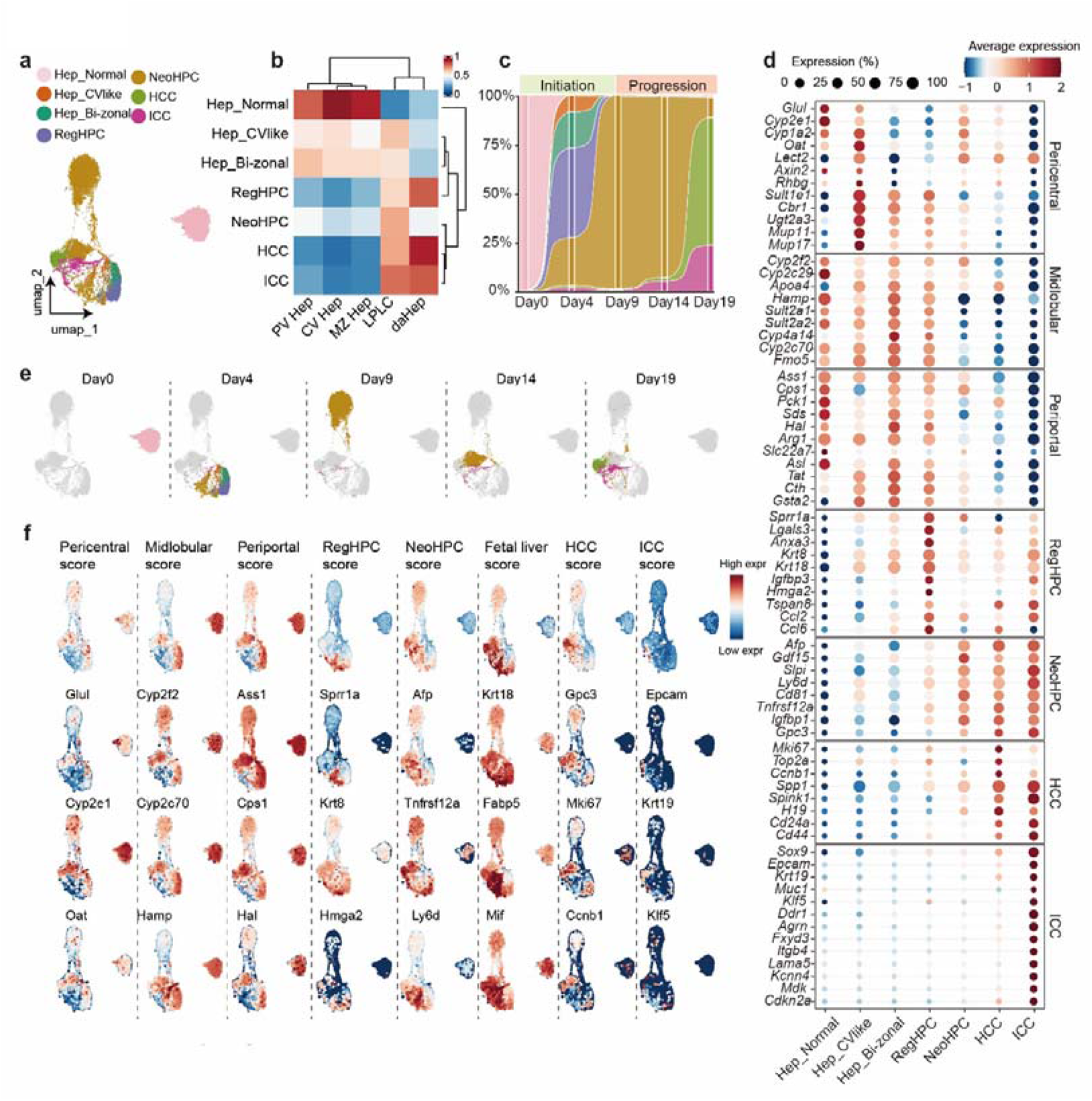
Hepatocytes traverse regenerative and premalignant states during liver tumor initiation. **(a)** UMAP embedding of the integrated single-cell transcriptomic data, color-coded by seven major identified cell states: normal hepatocytes (Hep_Normal), central vein-like hepatocytes (Hep_CVlike), bi-zonal hepatocytes (Hep_Bi-zonal), Regenerative hepatic progenitor cells (RegHPC), neoplastic hepatic progenitor cells (NeoHPC), hepatocellular carcinoma cells (HCC), and intrahepatic cholangiocarcinoma cells (ICC). **(b)** Heatmap showing the correlation scores mapping between the seven identified cell states (rows) and reference liver cell types (columns), including portal vein hepatocytes (PV Hep), central vein hepatocytes (CV Hep), mid-zone hepatocytes (MZ Hep), liver progenitor-like cells (LPLC), and disease-associated hepatocytes (daHep). The color gradient indicates the strength of the signature match. **(c)** Stacked area plot showing the proportions of the seven hepatocyte states across the indicated time points (Day 0, Day 4, Day 9, Day 14, and Day 19). The timeline is broadly partitioned into “Initiation” and “Progression” phases. **(d)** Bubble plot displaying the expression profiles of state-specific marker genes across the seven identified cell states (columns). Markers are grouped into pericentral, midlobular, and periportal zonal gene sets, followed by distinct transcriptional signatures specific to the RegHPC, NeoHPC, HCC, and ICC clusters. The color scale represents the scaled average expression level, and the dot size indicates the percentage of cells within each cluster expressing the respective gene. **(e)** Split UMAP plots showing the distribution of captured cells at each indicated time point (Day 0, Day 4, Day 9, Day 14, and Day 19). In each panel, cells from the corresponding time point are highlighted in color according to their state identities defined in (a), whereas cells from all other time points are displayed in light gray. **(f)** UMAP visualization of signature scores and representative gene expression profiles across the cell atlas. The top row displays the computed signature module scores for Pericentral, Midlobular, Periportal, RegHPC, NeoHPC, Fetal liver, HCC, and ICC states. The subsequent rows display expression levels of key representative marker genes corresponding to each respective cell state. Expression intensity and signature scores are color-coded by a blue-to-red gradient indicating low to high expression levels or signature enrichment.

We next annotated these hepatocyte states by integrating reference-based transcriptomic similarity with canonical marker-gene expression^21,72^. Hep_Normal cells showed the strongest similarity to homeostatic zonated hepatocytes, with high correlation coefficients with pericentral, midlobular, and periportal hepatocytes (CV Hep, r = 0.94; MZ Hep, r = 0.89; PV Hep, r = 0.80), and only weak similarity to liver progenitor-like cells ^72^(LPLCs; r = 0.19) or disease-associated hepatocytes^21^ (daHeps; r = 0.32) (Figure 2b; Supplementary Figure 3a, b; Supplementary Table 4). Consistently, Hep_Normal cells retained expression of mature hepatocyte and zonation-associated genes, supporting their identity as homeostatic hepatocytes^24,88^.

Among the oncogene-expressing hepatocyte-derived populations, Hep_CVlike cells retained partial pericentral identity, showing moderate similarity to CV hepatocytes (r = 0.57) and expressing canonical pericentral markers, including Glul, Cyp2e1, Oat, Lect2, and Rhbg^24,88–90^ (Figure 2b, d, f; Supplementary Table 4). However, these cells also showed increased similarity to injury-associated LPLCs^72^ (r = 0.64), suggesting that Hep_CVlike represents a pericentral-like hepatocyte state exposed to pathological stress rather than a fully homeostatic CV population^91^.

Hep_Bi-zonal cells displayed a distinct mixed zonation identity (Figure 2b, d, f; Supplementary Table 4). These cells showed comparable similarity to periportal and midlobular hepatocytes (PV Hep, r = 0.64; MZ Hep, r = 0.59), while expressing representative midlobular and periportal markers such as Cyp2f2, Cyp2c70, Ass1, Cps1, and Hal ^24,88,90^(Figure 2d). Hep_Bi-zonal cells also showed moderate similarity to LPLCs (r = 0.56), indicating that they preserve key metabolic features of non-pericentral hepatocytes while acquiring early injury-associated transcriptional features^27,30,72^.

RegHPC cells were distinguished by both reference-based and marker-based evidence of an injury-repair identity (Figure 2b, d, f; Supplementary Figure 3a, b). These cells showed strong similarity to LPLCs^92^ (r = 0.60) and even higher similarity to daHeps^21^ (r = 0.80), indicating convergence with hepatocyte states previously observed in chronic liver injury and fatty liver disease progression^21,72^. Consistently, RegHPC cells robustly expressed injury-response and tissue-repair-associated genes, including Sprr1a, Krt8, Lgals3, and Ly6d (Figure 2d). Previous studies have associated LPLCs with hepatocyte dedifferentiation and regenerative repair during liver injury^72,91,93^. Consistent with these observations, the coordinated expression of Sprr1a, Krt8, Lgals3, and Ly6d in RegHPC cells is indicative of an epithelial injury-response program accompanied by ductular-like remodeling and progenitor-associated regenerative features. These features support the annotation of RegHPC as a regenerative, injury-associated progenitor-like hepatocyte state^27,27,72,91,93–95^.

NeoHPC cells showed a related but more fetal-like and plastic transcriptional identity (Figure 2b, d, f; Supplementary Figure 3c, d). Compared with RegHPC, NeoHPC cells showed the highest similarity to LPLCs among the progenitor-like states (r = 0.69), while retaining appreciable similarity to daHeps (r = 0.48). In parallel, NeoHPC cells displayed markedly elevated fetal liver signature scores and increased expression of stemness- and oncofetal-associated genes, including Afp, Tnfrsf12a, Gpc3, and Fabp5 (Figure 2d). Afp and Gpc3 are well-established markers of fetal-like and oncofetal hepatocyte programs^96^, whereas Tnfrsf12a has been associated with injury-induced progenitor activation and regenerative plasticity^97^. These observations define NeoHPC as a fetal-like, highly plastic progenitor-like hepatocyte state.

The malignant populations were also transcriptionally aligned with disease-associated hepatocyte programs. HCC showed strong similarity to daHeps (r = 0.89) and LPLCs (r = 0.69), whereas ICC showed high similarity to both LPLCs (r = 0.77) and daHeps (r = 0.81) (Figure 2b; Supplementary Figure 3a, b). Together with their marker profiles, these correlations indicate that both malignant populations retain disease-associated transcriptional features rather than resembling homeostatic hepatocytes.

We then examined the temporal distribution of these hepatocyte states during tumor initiation and progression. The relative abundance of each population changed progressively across time points, revealing extensive remodeling of the hepatocyte compartment (Figure 2c, e). Early stages were characterized by pronounced heterogeneity, including Hep_CVlike, Hep_Bi-zonal, RegHPC, and NeoHPC states, whereas later stages were increasingly enriched for NeoHPC and malignant populations (Figure 2c, e). These temporal patterns indicate that liver tumor initiation is accompanied by a progressive expansion of disease-associated hepatocyte states.

Collectively, these findings demonstrate that hepatocyte transformation proceeds through sequential regenerative and premalignant progenitor states rather than a direct transition from homeostatic hepatocytes to malignancy.

### Lineage reconstruction identifies Hep_Bi-zonal cells as the cellular origin of liver tumor initiation

Although transcriptomic analyses revealed a continuum of disease-associated hepatocyte states, it remained unclear whether these populations represented bona fide lineage intermediates or alternative cell-fate trajectories. To resolve their lineage relationships, we leveraged the accumulated editing scars generated by CERAMIC to reconstruct clonal phylogenies at single-cell resolution.

Before phylogenetic reconstruction, we first evaluated the recovery efficiency and integrity of CERAMIC lineage barcodes across all hepatocyte-derived populations and developmental stages. Recovery of informative lineage barcodes remained consistently high throughout tumor initiation, with 98.5-100% of cells across Hep_CVlike, Hep_Bi-zonal, RegHPC, NeoHPC, HCC, and ICC populations retaining informative edited lineage barcodes (Figure 3a; Supplementary Table 5). Single-cell allele calling and phylogenetic reconstruction were performed as described in Methods. Individual cells retained an average of 2.23-4.35 edited recording units, with the greatest recording depth observed in Hep_CVlike and Hep_Bi-zonal cells, while fully edited recording arrays remained detectable across all cell states (Figure 3a). Consistent with this high recovery efficiency, informative lineage barcodes were recovered from 3,282, 5,000, 2,570, and 2,113 cells at Day 4, Day 9, Day 14, and Day 19, respectively, providing sufficient lineage information for robust phylogenetic reconstruction (Supplementary Figure 4 b, f, j, n). Moreover, recovered editing scars were predominantly confined to individual recording modules, with single-target insertions and deletions accounting for the vast majority of editing events across all time points, whereas inter-site deletions spanning more than three recording modules remained rare during early tumor initiation (0.1% at Day 4, 19.1% at Day 9, and 9.1% at Day 14), increasing only after prolonged recording at Day 19 (33.9%), consistent with the cumulative nature of continuous CRISPR editing (Supplementary Figure 4 a, e, i, m). In addition, each recording site generated a rich repertoire of distinct editing outcomes, with individual sgRNA target sites producing approximately 100-300 unique alleles across the dataset (Supplementary Figure 4c, g, k, o). This extensive editing diversity substantially expanded the lineage information encoded at each recording module, thereby increasing the resolving power for phylogenetic reconstruction. Together, these results demonstrate that CERAMIC continuously generates stable, highly diverse, and information-rich lineage barcodes while preserving recording-module integrity throughout liver tumor initiation, thereby enabling accurate single-cell phylogenetic reconstruction of hepatocyte fate transitions (Supplementary Figure 4d, h, l, p).

**Figure 3.**
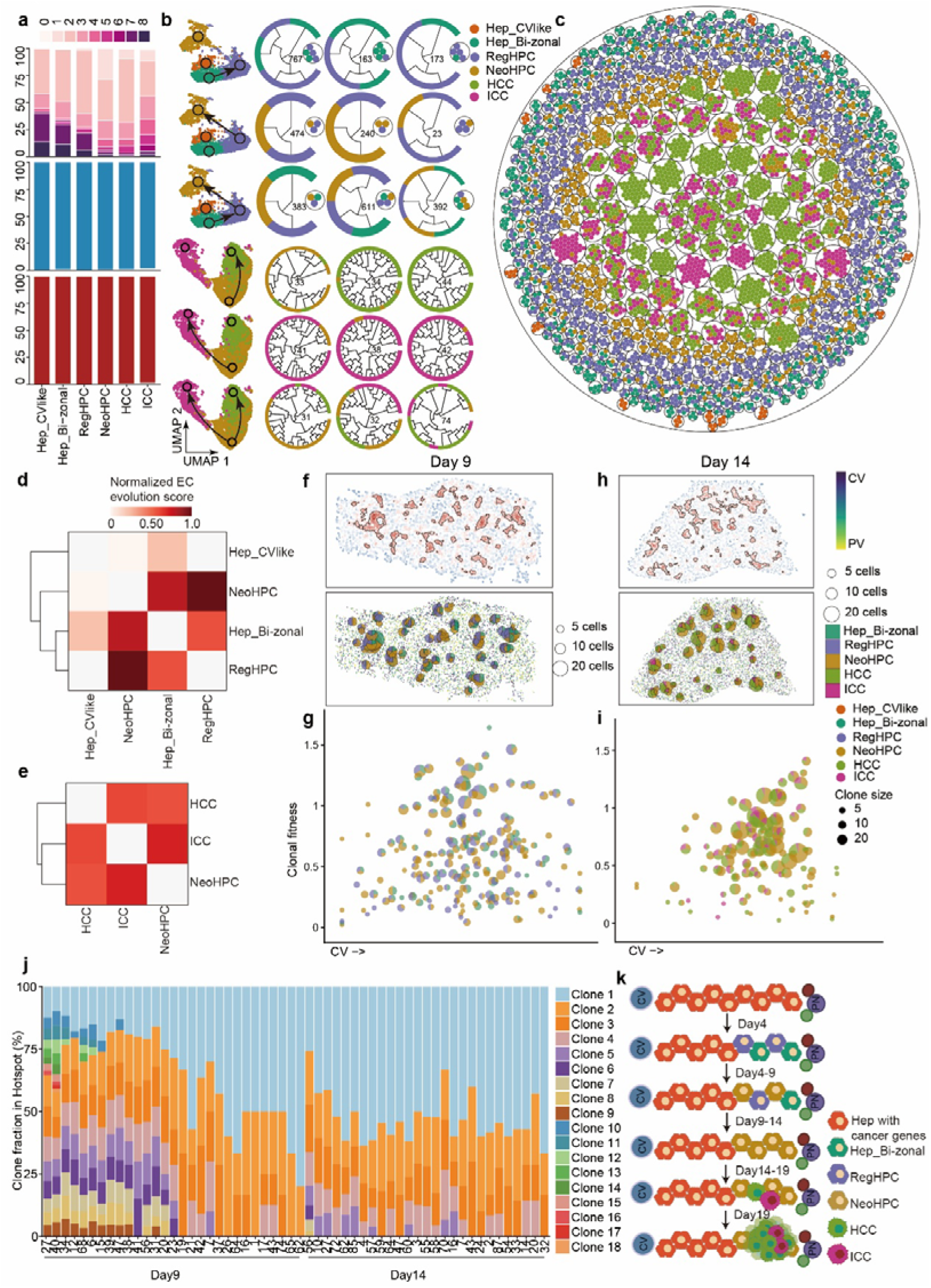
High-resolution lineage reconstruction and spatial clonal tracking identify Hep_Bi-zonal cells as the primary tumor-fated hepatocyte state. **(a)** Quantitative assessment of CERAMIC performance across the six indicated hepatocyte-derived cell states (horizontal axis). Top, stacked bar chart showing the distribution of recovered target-array (TA) module counts (0-8; pink-to-purple gradient). Middle, detection efficiency of lineage barcodes within the valid cell population. Bottom, proportions of edited and unedited barcodes. **(b)** Representative phylogenetic trees of individual reconstructed clones displayed together with the cell-state UMAP embedding. The left panels show the corresponding clones on the UMAP embedding, with black arrows indicating the highlighted branches. The middle and right panels show circular phylogenetic trees containing Hep_CVlike, Hep_Bi-zonal, RegHPC, NeoHPC, HCC, and ICC cells. Inner numbers indicate clone IDs, and the outer colored rings denote the corresponding cell states. **(c)** Circle-packing visualization of reconstructed clones. Each large circle represents an individual clone, and each dot represents a single cell colored according to its annotated cell state. **(d, e)** Hierarchical clustering and heatmaps of normalized evolutionary coupling (EC) scores. Color indicates normalized EC scores (0-1.0). Panel **(d)** shows EC scores among Hep_CVlike, Hep_Bi-zonal, RegHPC, and NeoHPC states, and panel **(e)** shows EC scores among NeoHPC, HCC, and ICC states. **(f, g)** Spatial clonal analysis at Day 9. **(f)** Top, spatial tissue coordinates with tumor density contours (red outlines). Bottom, spatial distribution of reconstructed clones overlaid on liver zonation coordinates (CV-to-PV axis). Pie charts represent individual clones; pie size indicates clone size, and pie sectors indicate the proportions of different cell states within each clone. **(g)** Clonal fitness plotted against spatial position along the CV-to-PV axis. The y-axis shows the clonal fitness score, the x-axis shows spatial position along the liver zonation axis, and bubble color indicates the dominant cell state. **(h, i)** Spatial clonal analysis at Day 14. **(h)** Top, spatial tissue coordinates with tumor density contours (red outlines). Bottom, spatial distribution of reconstructed clones overlaid on liver zonation coordinates (CV-to-PV axis). Pie charts represent individual clones; pie size indicates clone size, and pie sectors indicate the proportions of different cell states within each clone. **(i)** Clonal fitness plotted against spatial position along the CV-to-PV axis. The y-axis shows the clonal fitness score, the x-axis shows spatial position along the liver zonation axis, and bubble color indicates the dominant cell state. **(j)** Stacked bar chart showing the clonal composition of individual spatial tumor hotspots at Day 9 and Day 14. The y-axis indicates clone fraction, each column represents one hotspot, and colors denote individual clones (Clone 1-18). **(k)** Schematic illustrating hepatocyte states and reconstructed lineage relationships across the indicated time points (Day 4-Day 19). Colors correspond to the hepatocyte states shown throughout the study. PN = portal node (a vasculature structure composed from the portal vein (purple), hepatic artery (red) and bile duct (green)), CV = central vein (blue).

We next reconstructed representative single-cell lineage phylogenies from early-stage hepatocyte populations to resolve the lineage origins of the intermediate progenitor states (Figure 3b; Supplementary Table 6; see Methods). This analysis included 4,379 Hep_Normal cells, 275 Hep_CVlike cells, 607 Hep_Bi-zonal cells, 1,499 RegHPC cells, 8,294 NeoHPC cells, 1,479 HCC cells, and 749 ICC cells. Lineage clones were identified based on patristic distances within reconstructed phylogenetic trees (see Methods). Across independently reconstructed lineage trees, Hep_Bi-zonal cells consistently occupied ancestral positions within lineage trees and gave rise to RegHPC populations, whereas Hep_CVlike cells showed little contribution to this transition. RegHPC cells subsequently generated NeoHPC cells, which frequently served as immediate ancestral intermediates for both HCC and ICC lineages, revealing a hierarchical lineage trajectory from Hep_Bi-zonal through RegHPC and NeoHPC to malignant cell fates (Figure 3b). Together, these lineage relationships demonstrate that the hepatocyte state continuum inferred from transcriptomic analyses reflects a bona fide lineage hierarchy linking Hep_Bi-zonal cells to malignant cell fates, rather than simply representing transcriptionally similar but lineage-unrelated cell states.

To globally visualize clonal relationships, we next reconstructed the overall clonal architecture of hepatocyte-derived populations (Figure 3c; Supplementary Table 6). Consistent with the representative phylogenies, RegHPC and NeoHPC cells were predominantly found within clones containing Hep_Bi-zonal cells. Quantitatively, among filtered clones, 250 of 293 Hep_Bi-zonal-containing branches progressed into RegHPC or NeoHPC states (85.3%), whereas none of the 16 Hep_CVlike-containing branches did so (0%; Fisher’s exact test, P = 4.93 × 10 ¹³). This strong asymmetry remained evident under a more stringent endpoint definition requiring NeoHPC-containing clones, with 131 of 293 Hep_Bi-zonal-containing branches reaching the NeoHPC state compared with 0 of 16 Hep_CVlike-containing branches. In contrast, Hep_CVlike cells largely formed isolated clones and only rarely shared common ancestry with downstream progenitor or malignant populations. These results provide quantitative support for the conclusion that Hep_Bi-zonal cells, rather than Hep_CVlike cells, preferentially seed the tumor-fated progenitor trajectory.

Building on the reconstructed phylogenies, we next quantified lineage relationships among hepatocyte states, we calculated normalized evolutionary coupling scores based on reconstructed phylogenetic trees (Figure 3d, e; See Methods). Hep_Bi-zonal cells exhibited strong evolutionary coupling with both RegHPC and NeoHPC populations, further supporting the conclusion that these intermediate states originate from the Hep_Bi-zonal compartment. Likewise, NeoHPC cells exhibited the highest evolutionary coupling scores with both HCC and ICC populations, indicating that NeoHPC represents the immediate progenitor state from which malignant diversification emerges (Figure 3d, e). Together, these analyses establish a hierarchical lineage hierarchy in which Hep_Bi-zonal cells progress through regenerative intermediates before entering a highly plastic NeoHPC state that subsequently diversifies into both HCC and ICC.

We next asked whether these lineage-defined states exhibit distinct spatial organization within the liver lobule. To address this question, we mapped lineage information with spatial transcriptomic datasets to investigate the anatomical distribution of lineage-defined clones (Figure 3f, g; Supplementary Table 6). We first identified actively expanding tumor hotspots from local tumor-cell density maps and subsequently reconstructed hotspot-specific lineage phylogenies to define individual lineage clones (see Methods). Systematic quality assessment demonstrated that more than 60% of tumor-associated cells within these regions retained informative edited barcodes, enabling reliable spatial lineage reconstruction (Supplementary Figure 3r-z).

Mapping reconstructed lineage clones onto tissue coordinates revealed a striking spatial pattern, with expanded clones consistently enriched along the midlobular-to-periportal axis rather than the pericentral region (Figure 3f, g; Supplementary Table 6). Notably, this positional bias was observed in both early progenitor-derived clones and later malignant clones undergoing extensive expansion. Moreover, clone fitness, estimated using a previously described phylogeny-based fitness inference framework (see Methods)^55^, showed a progressive increase toward periportal regions, particularly at Day 14, indicating that tumor-initiating clones preferentially expanded along the midlobular-periportal axis (Figure 3f-i). This spatial bias suggests that regional features associated with these lobular zones, including intrinsic metabolic programs and local microenvironmental cues, may collectively create conditions that favor the emergence and expansion of tumor-initiating cells.

Finally, we characterized how clonal composition changed within individual tumor hotspots as lesions progressed (Figure 3j; Supplementary Figure 5). At Day 9, hotspots were highly polyclonal, comprising numerous intermingled lineage-defined clones with no single clone predominating (Figure 3j; Supplementary Figure 5a, c, e; Supplementary Table 6). Quantification of effective clonal diversity, calculated as the exponential of Shannon entropy (see Methods), showed that Day 9 hotspots contained 6.18 ± 3.91 effective clones (Supplementary Figure 5a). By Day 14, effective clonal diversity had significantly contracted to 3.33 ± 0.82 effective clones (P = 0.0217), indicating progressive loss of clonal diversity during tumor expansion (Figure 3j; Supplementary Figure 5b, d, f; Supplementary Table 6). In parallel, the dominant clone ratio (see Methods) increased from 34.1 ± 20.0% at Day 9 to 52.8 ± 9.7% at Day 14 (P = 1.76 × 10), demonstrating the emergence of dominant high-fitness lineages (Supplementary Figure 5a, b). Spatial clone maps and hotspot-associated phylogenies further supported this transition: Day 9 lesions contained many small, intermixed clones, whereas Day 14 lesions were increasingly organized around fewer, larger, and more dominant lineage-defined clones (Supplementary Figure 5 c-f). Thus, these findings indicate that liver tumor initiation proceeds through a transition from polyclonal seeding to oligoclonal outgrowth, reflecting strong clonal competition and an evolutionary bottleneck in which only a subset of clones sustains continued tumor expansion.

Collectively, these lineage-tracing analyses reveal that Hep_Bi-zonal cells constitute the principal lineage origin of tumor-initiating progenitors during liver tumorigenesis. Through a hierarchical progression involving RegHPC and NeoHPC intermediates, these cells ultimately generate both HCC and ICC lineages. In parallel, spatially resolved lineage reconstruction demonstrates that this evolutionary process is preferentially localized to the midlobular-to-periportal niche and is accompanied by progressive clonal selection, ultimately leading to oligoclonal tumor expansion.

### Transcriptional and regulatory programs distinguish tumor-fated hepatocytes from failed-to-transform lineages

Although lineage tracing established Hep_Bi-zonal cells as the principal source of tumor-fated progenitors with lineage plasticity, the molecular programs that enable this transition remained unclear (Figure 4). We therefore sought to identify the molecular programs and key regulatory factors that drive hepatocyte fate transitions during liver tumor initiation by systematically interrogating transcriptional states, gene regulatory networks, and spatial signaling landscapes.

**Figure 4.**
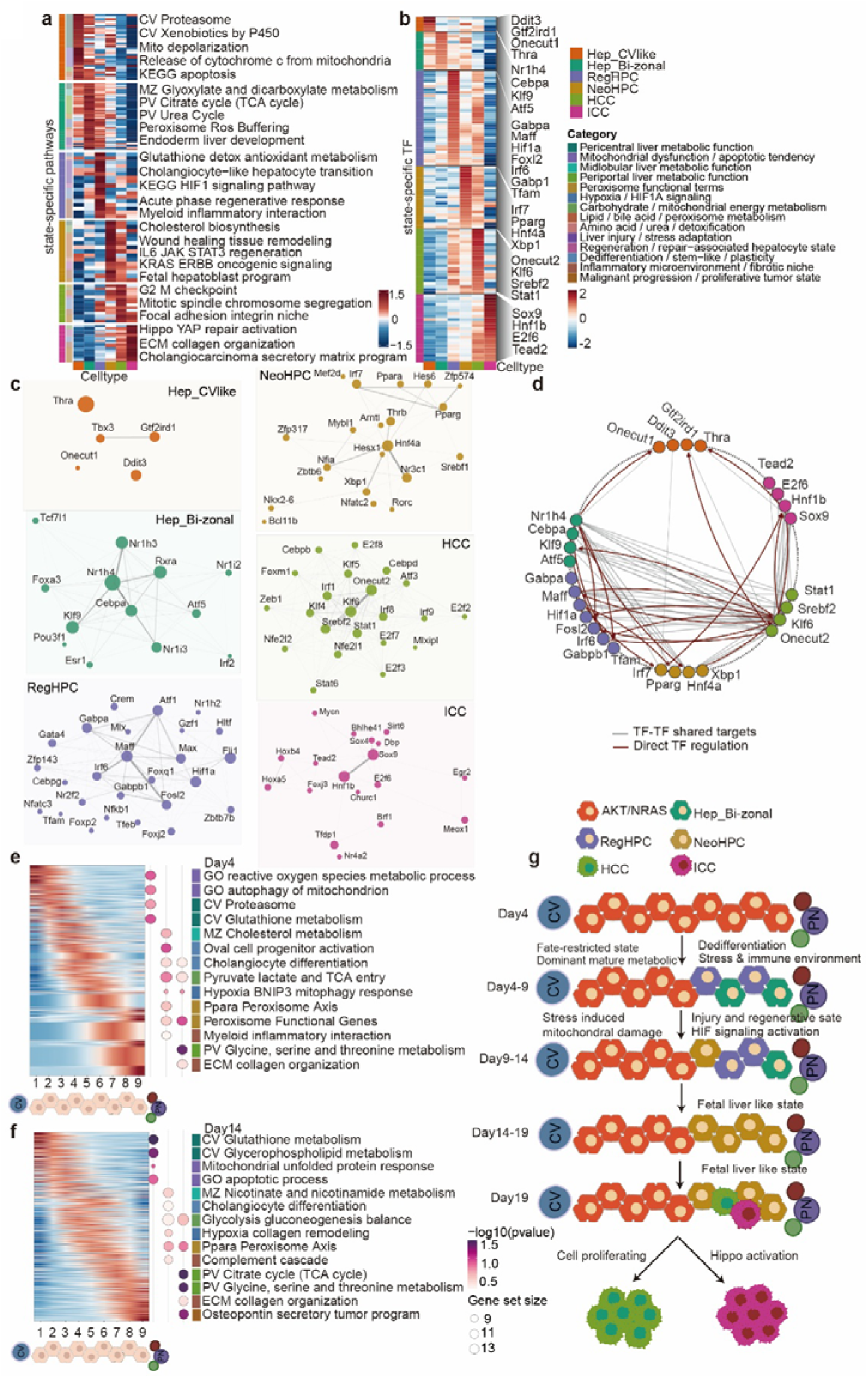
Regulatory and metabolic programs distinguish tumor-fated Hep_Bi-zonal cells from Hep_CVlike cells. **(a)** Heatmap showing pathway module scores across Hep_CVlike, Hep_Bi-zonal, RegHPC, NeoHPC, HCC, and ICC cell states. Pathway scores were calculated at the single-cell level from predefined gene signatures, averaged within each cell state, and standardized for each pathway as row-wise z-scores. Red indicates relatively higher pathway scores and blue indicates relatively lower pathway scores (clipped to −1.5 to 1.5). Rows represent pathway modules grouped into liver zonation-associated metabolism, mitochondrial function, apoptosis, peroxisome function, hypoxia/HIF1A signaling, liver injury and stress response, regeneration, cellular plasticity, inflammatory and fibrotic signaling, and malignant-associated programs. Columns represent the indicated hepatocyte states. Row annotations indicate pathway categories. **(b)** Heatmap showing the activity of selected transcription factor (TF) regulons across Hep_CVlike, Hep_Bi-zonal, RegHPC, NeoHPC, HCC, and ICC cell states. Regulon activity was quantified using pySCENIC AUCell scores, averaged within each cell state, and displayed as row-wise z-scored regulon AUC values. TFs are grouped according to their annotated cell-state specificity, and columns represent the indicated cell states. Red indicates relatively higher regulon activity and blue indicates relatively lower regulon activity. **(c)** Co-target networks of selected transcription factors for each cell state. Individual panels correspond to Hep_CVlike, Hep_Bi-zonal, RegHPC, NeoHPC, HCC, and ICC. Nodes represent TFs, with node colors indicating cell-state identity and node sizes indicating the number of shared downstream target genes. Edges represent TF-TF co-target relationships defined by the overlap of downstream target genes between pySCENIC regulons, and edge width is proportional to the degree of target overlap. Node positions were determined according to network connectivity. **(d)** Network showing the relationships among selected hub transcription factors across Hep_CVlike, Hep_Bi-zonal, RegHPC, NeoHPC, HCC, and ICC cell states. Nodes represent hub TFs, with node colors indicating the corresponding cell state and node sizes indicating network connectivity. Gray edges represent TF-TF co-target relationships based on shared downstream target genes, and dark red directed edges represent regulatory relationships inferred from pySCENIC regulons. Nodes are arranged according to cell-state annotation in a circular layout. **(e, f)** Spatial pathway analysis across nine concentric spatial layers at Day 4 (e) and Day 14 (f). Spatial transcriptomic cells were grouped into nine layers according to their relative spatial position, with layers 1-3 corresponding to the pericentral region, layers 4-6 to the midlobular region, and layers 7-9 to the periportal region. Left, heatmaps showing scaled expression of genes ordered according to their spatial layer preference. Rows represent selected genes and columns represent the mean expression of each spatial layer. Red indicates relatively higher expression and blue indicates relatively lower expression. Right, dot plots showing representative enriched GO and KEGG pathways for the pericentral, midlobular, and periportal layer groups. Dot size indicates the number of genes in each pathway, and dot color indicates enrichment significance (-log₁₀ *P* value). Pathways are grouped according to the functional categories shown in (a). **(g)** Schematic illustrating hepatocyte states, spatial distribution, and pathway annotations across the indicated time points (Day 4-Day 19). Colors correspond to the hepatocyte states shown throughout the study. PN = portal node (a vasculature structure composed from the portal vein (purple), hepatic artery (red) and bile duct (green)), CV = central vein (blue).

We first compared the two earliest divergent hepatocyte populations, Hep_CVlike and Hep_Bi-zonal, to identify the transcriptional programs associated with their distinct lineage fates (Figure 4a; Supplementary Table 7). The non-tumorigenic Hep_CVlike population predominantly retained canonical pericentral liver programs, including xenobiotic metabolism, cytochrome P450-associated pathways, and proteostasis-associated pathways, consistent with the characteristic metabolic specialization of pericentral hepatocytes^24,26^(Figure 4a; Supplementary Figure 6a). Notably, these cells also showed high scores for mitochondrial dysfunction and cell-death-associated programs, including mitochondrial depolarization, cytochrome c release, and apoptotic signaling. Together, these observations suggest that Hep_CVlike cells experience substantial metabolic and mitochondrial stress in the fatty liver-associated pathological environment and are consequently biased toward a non-tumorigenic fate.

In contrast, the tumor-fated Hep_Bi-zonal population maintained characteristic midlobular and periportal metabolic programs, including glyoxylate and dicarboxylate metabolism, citrate cycle activity, and urea-cycle-associated functions (Figure 4a; Supplementary Table 7). Importantly, these cells also displayed higher scores for peroxisome-associated, reactive oxygen species-buffering, and endodermal liver-development programs. These findings suggest that Hep_Bi-zonal cells preserve a metabolically adaptive state that may facilitate tolerance to fatty liver-associated lipotoxic and oxidative stress while maintaining lineage plasticity. Together, these metabolic and developmental programs are consistent with a cell state capable of supporting survival, regenerative remodeling, and tumor-fated transition^98^.

As hepatocytes progressively exited their mature identity, the intermediate RegHPC population acquired a transcriptional profile characteristic of tissue injury and regeneration (Figure 4a; Supplementary Table 7). Compared with mature hepatocytes, RegHPC cells displayed higher scores for glutathione-mediated antioxidant programs, hepatocyte-to-cholangiocyte transition signatures, acute-phase regenerative responses, hypoxia-responsive gene sets, and myeloid inflammatory interaction programs. These transcriptional features are consistent with previous studies describing regenerative hepatocyte populations that emerge during chronic liver injury and tissue repair^27,30,72,93,99,100^. Consistent with this interpretation, published injury-, stress-, and regeneration-associated gene signatures^72^ became prominently represented in RegHPC cells, further supporting the view that RegHPC corresponds to a transient regenerative intermediate linking mature hepatocytes to neoplastic progenitor cells (Supplementary Figure 6f).

Subsequently, NeoHPC cells acquired a distinct fetal-like progenitor state characterized by enhanced developmental plasticity and acquisition tumor-associated transcriptional programs (Figure 4a; Supplementary Table 7). Compared with RegHPC cells, NeoHPCs displayed higher scores for cholesterol biosynthesis, wound-healing and tissue-remodeling programs, IL6-JAK-STAT3 signaling, KRAS-responsive programs, and fetal hepatoblast signatures. These transcriptional features are consistent with previous studies showing that liver tumorigenesis is accompanied by reactivation of developmental and fetal-like programs associated with cellular plasticity and malignant progression^31,32,101,102^.

Following acquisition of this highly plastic progenitor state, the two malignant lineages underwent further lineage-specific specialization (Figure 4a; Supplementary Table 7). HCC cells preferentially displayed higher scores for proliferative programs, including G2/M checkpoint regulation, mitotic spindle organization, and chromosome segregation, consistent with active cell-cycle progression. In contrast, ICC cells exhibited higher scores for Hippo-YAP-associated programst^100,103,104^., extracellular matrix organization^105^, collagen remodeling^106^, focal adhesion, and cholangiocarcinoma secretory matrix signatures, consistent with progressive biliary lineage commitment together with extensive remodeling of the surrounding microenvironment. Collectively, these findings suggest that liver tumor initiation is not simply accompanied by progressive dedifferentiation. Instead, hepatocytes undergo a highly ordered sequence of transcriptional state transitions, progressing from metabolic adaptation, to an injury-adaptive regenerative intermediate, to a highly plastic oncofetal progenitor state, before ultimately diverging into lineage-specialized HCC and ICC cells.

To determine whether transcriptome-based trajectory inference recapitulated the lineage relationships identified by CERAMIC, we reconstructed pseudotime trajectories across hepatocyte populations. Early pseudotime trajectories recapitulated a progressive transition from Hep_Bi-zonal cells through RegHPC to NeoHPC states, accompanied by dynamic remodeling of regenerative, metabolic, and developmental programs (Supplementary Figure 6c-f; Supplementary Table 8; see Methods). Similarly, late-stage pseudotime trajectories diverged from NeoHPC toward HCC and ICC lineages, accompanied by progressive representation of HCC-associated proliferative programs and ICC-associated biliary differentiation and matrix-remodeling programs (Supplementary Figure 6g-j; Supplementary Table 8). Consistent with these lineage trajectories, single-cell analyses revealed progressive enrichment of transcriptional programs associated with tissue injury, hypoxia, inflammation, cellular senescence, and fetal liver identity along the Hep_Bi-zonal, RegHPC, NeoHPC continuum, with the highest activity observed in tumor-fated cell states during tumor progression.

We therefore reconstructed cell-state-specific regulon networks using pySCENIC^107,108^ to identify the upstream transcription factors governing these molecular programs. (Figure 4b, c; Supplementary Table 7; see Methods). Distinct TF modules were associated with each stage of tumor evolution. Hep_CVlike cells were predominantly associated with stress- and differentiation-related regulons, including Ddit3, Gtf2ird1, Onecut1, and Thra, consistent with their high mitochondrial stress and apoptotic program scores. In contrast, Hep_Bi-zonal cells were associated with metabolic and hepatocyte-identity regulators, including Nr1h4 (FXR)^109,110^, Cebpa, Klf9, and Atf5^111,112^, supporting maintenance of metabolic homeostasis and adaptive responses to fatty liver-associated stress. Transition into the RegHPC state was accompanied by a regulatory module centered on Gabpa, Maff, Hif1a, Fosl2, Irf6, Gabpb1, and Tfam, consistent with coordinated hypoxia adaptation^113,114^, mitochondrial remodeling^115,116^, and inflammatory signaling^117,118^. NeoHPC cells acquired a distinct transcriptional circuitry dominated by Irf7, Pparg, Hnf4a, and Xbp1, regulators associated with stress adaptation, metabolic remodeling, inflammatory signaling, and cellular stress adaptation^103,119–124^. Finally, lineage-specific malignant programs emerged, with Onecut2, Klf6, Srebf2, and Stat1 constituting major HCC-associated regulons, whereas Sox9^125^, Hnf1b, E2f6, and Tead2 were preferentially associated with ICC cells.

We further examined the relationships among these core regulators by constructing an integrated TF-TF co-regulatory network (Figure 4d; Supplementary Figure6b; Supplementary Table 7; See Methods). Notably, TFs defining the Hep_Bi-zonal state shared extensive downstream target genes with the core RegHPC regulatory module, including approximately 84 shared target genes. Moreover, several Hep_Bi-zonal regulators displayed direct regulatory links to key RegHPC-associated TFs, involving 3 inferred regulatory edges. This regulatory architecture suggests that Hep_Bi-zonal cells are transcriptionally primed for transition into the regenerative progenitor state, rather than representing simply another mature hepatocyte population. In contrast, the Hep_CVlike regulatory network showed comparatively limited overlap with the RegHPC program, with only 3 shared target genes, consistent with lineage-tracing evidence demonstrating that Hep_CVlike cells rarely contribute to downstream tumor-fated populations.

To determine whether these transcriptional programs exhibited corresponding spatial organization within the liver lobule, we next quantified spatial gene-set scores along the central vein (CV)-to-portal vein (PV) axis using spatial transcriptomic data (Figure 4e, f; Supplementary Figure 7; Supplementary Table 7). At Day 4, pericentral regions showed higher scores for mitochondrial stress and damage-associated programs, including mitophagy, mitochondrial autophagy, proteasome activity, and glutathione metabolism. In contrast, the midlobular-periportal axis displayed higher scores for peroxisome-associated programs, PPAR signaling, inflammatory interaction signatures, progenitor-cell activation programs, and cholangiocyte differentiation-associated gene sets (Figure 4e). These spatial patterns closely mirrored the transcriptional differences observed between Hep_CVlike and Hep_Bi-zonal cells at single-cell resolution.

As tumor progression advanced to Day 14, the midlobular-periportal region increasingly acquired features of a tumor-permissive spatial niche (Figure 4f; Supplementary Figure 7). This domain showed higher scores for complement activation, extracellular matrix organization, collagen remodeling, hypoxia-associated signaling, and osteopontin-secretory tumor programs, while maintaining cholangiocyte differentiation and metabolic adaptation signatures. Together, these observations indicate that the same spatial regions supporting the emergence of tumor-fated hepatocytes subsequently evolve into microenvironments associated with malignant expansion.

Taken together, these findings support a model in which hepatocytes exposed to the fatty liver-associated pathological environment adopt divergent molecular responses according to their underlying metabolic and regulatory states. Pericentral-like hepatocytes preferentially show mitochondrial dysfunction and apoptotic programs, resulting in a non-tumorigenic fate (Figure 4g). In contrast, hepatocytes residing along the midlobular-periportal axis retain metabolic programs associated with peroxisomal metabolism and antioxidant defense, acquire regenerative and hypoxia-associated transcriptional programs, and subsequently transition through RegHPC and NeoHPC states to generate malignant HCC and ICC lineages. This molecular framework helps explain why tumor-initiating potential is selectively acquired by hepatocytes residing in the midlobular--periportal region during fatty liver-associated hepatocarcinogenesis.

### Peroxisomal metabolism is required for progenitor-state formation and liver tumor initiation

Having identified Hep_Bi-zonal cells as the principal tumor-fated hepatocyte population, we next sought to determine which metabolic adaptations distinguish these cells from fate-restricted Hep_CVlike hepatocytes. Although both populations arise within the same fatty liver-associated pathological environment, Hep_Bi-zonal and Hep_CVlike ultimately adopt markedly different developmental trajectories, suggesting that their distinct cellular fates are associated with differential metabolic responses to chronic lipotoxic and inflammatory stress. We therefore systematically compared the transcriptional programs enriched between these two early hepatocyte states (Figure 5a; Supplementary Table 9).

**Figure 5.**
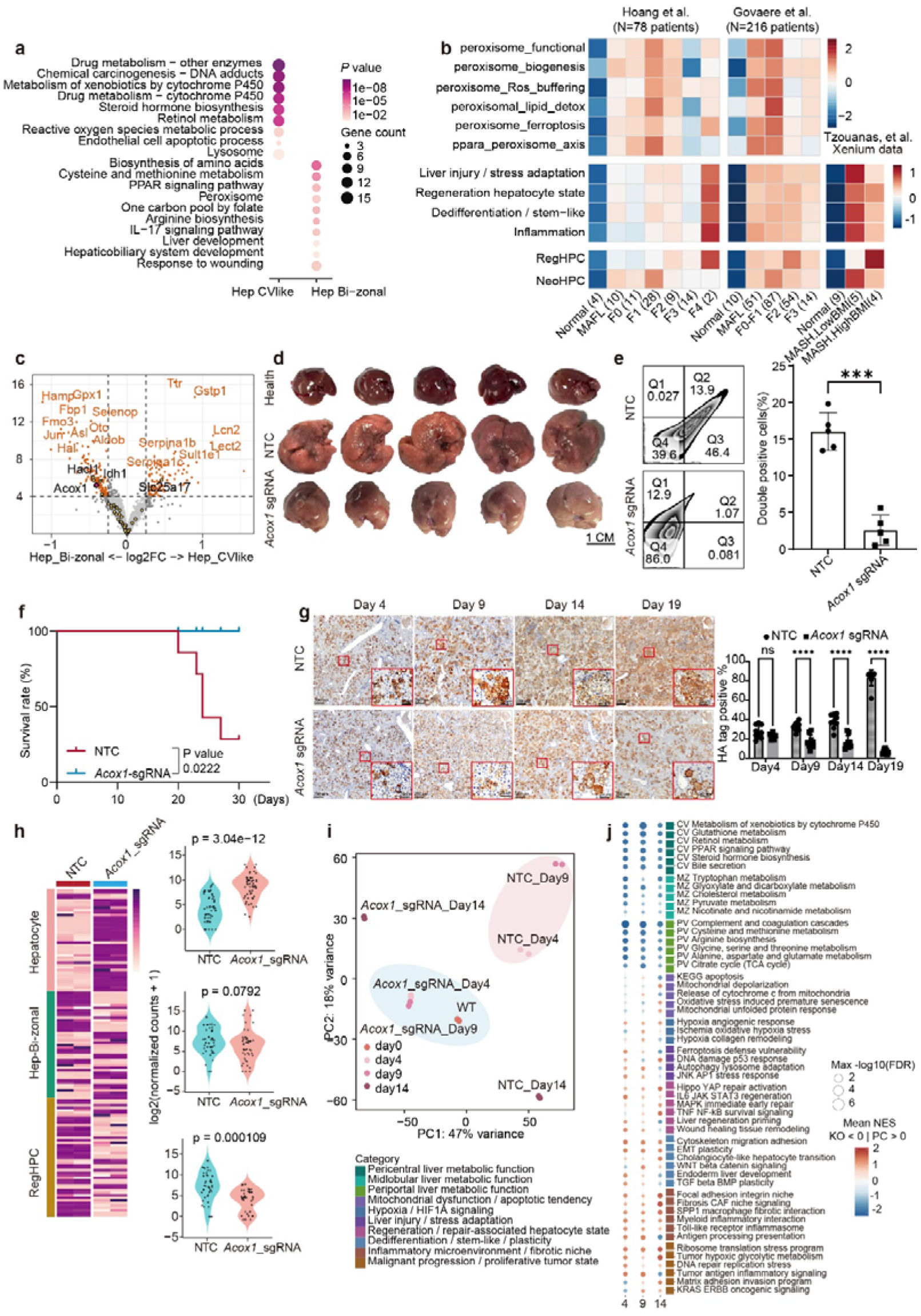
ACOX1-dependent peroxisomal metabolism is required for liver tumor initiation. **(a)** Dot plot showing enriched pathways for Hep_CVlike and Hep_Bi-zonal cells. Dot size indicates the number of genes associated with each pathway, and dot color indicates the *P* value. **(b)** Heatmaps showing pathway module scores across human liver disease datasets. Left, module scores across disease stages in the Hoang *et al.* cohort (n = 78). Middle, module scores across disease stages in the Govaere *et al.* cohort (n = 216). Right, module scores in the Xenium spatial transcriptomic dataset (Tzouanas *et al.*). Values are displayed as standardized z-scores. **(c)** Volcano plot showing differentially expressed genes between Hep_Bi-zonal and Hep_CVlike cells. Peroxisome-associated genes are highlighted in yellow, significantly differentially expressed genes in orange, and *Acox1* in pink. The horizontal dashed line indicates the significance threshold, and the x-axis shows the log₂ fold change. **(d)** Representative gross liver images from healthy control, non-targeting control (NTC), and Acox1 sgRNA-treated mice at Day 19 after induction. Scale bar, 1 cm. **(e)** Flow cytometric analysis of EGFP⁺tdTomato⁺ cells following Acox1 perturbation. Left, representative gating strategy for NTC and Acox1 sgRNA groups. Right, quantification of EGFP⁺tdTomato⁺ cells. Data were analyzed using a two-tailed unpaired *t* test with Welch’s correction (n = 5 mice per group; \*\*\**P* < 0.0001). **(f)** Kaplan-Meier survival curves for NTC and Acox1 sgRNA-treated mice during the 30-day observation period. *P* values were calculated using the log-rank Mantel-Cox test. **(g)** Immunohistochemical analysis of HA-tag-positive cells at Days 4, 9, 14, and 19 following *Acox1* perturbation. HA immunostaining detects HA-tagged AKT-expressing cells. Left, representative HA immunohistochemical staining (scale bars, 200 μm; enlarged views, 20 μm). Right, quantification of HA-positive cells. Data were analyzed by two-way ANOVA with Sidak’s multiple-comparisons test (n = 9 fields of view per group per time point; Day 4, adjusted *P* = 0.4507, ns; Day 9, adjusted *P* < 0.0001 (); Day 14, adjusted *P* < 0.0001 (); Day 19, adjusted *P* < 0.0001 (****)). **(h)** Bulk RNA-seq analysis of marker gene expression following Acox1 perturbation. Left, heatmap showing scaled expression of marker genes representing hepatocyte, Hep_Bi-zonal, and RegHPC states in NTC and Acox1 sgRNA samples. Right, violin plots showing normalized expression values (log₂(normalized counts + 1)) for marker genes in each gene set. Each dot represents one marker gene, and exact *P* values are indicated. **(i)** Principal component analysis (PCA) of bulk RNA-seq profiles from NTC and Acox1 sgRNA-treated samples collected at Days 0, 4, 9, and 14. **(j)** Dot plot showing pathway enrichment analysis of bulk RNA-seq data at Days 4, 9, and 14 following Acox1 perturbation. Pathways are grouped according to biological categories. Dot size indicates enrichment significance (-log₁₀ FDR), and dot color indicates the normalized enrichment score (NES).

Comparative pathway analysis revealed a pronounced metabolic divergence between the two populations (Figure 5a; Supplementary Table 9). Hep_CVlike cells largely retained canonical pericentral metabolic functions, including xenobiotic metabolism, cytochrome P450 activity, steroid metabolism and retinol metabolism^24,26,126^, while simultaneously exhibiting elevated mitochondrial depolarization, cytochrome-c release, oxidative stress and apoptotic programs. These observations are highly consistent with the transcriptional characteristics identified in Figure 4 and indicate that Hep_CVlike cells preserve a mature pericentral metabolic identity despite mounting evidence of progressive mitochondrial dysfunction and apoptotic stress. In contrast, Hep_Bi-zonal cells adopted a markedly more adaptive transcriptional state. Besides maintaining characteristic midlobular and periportal metabolic functions, these cells displayed coordinated activation of peroxisome-associated pathways, PPAR signaling, IL-17 inflammatory signaling, liver developmental programs and wound-response pathways. Peroxisomes and PPAR-regulated transcription are central to hepatic fatty-acid oxidation, lipid detoxification, and cellular redox homeostasis^127–129^, providing a potential framework through which these programs could support adaptation to lipotoxic and oxidative stress. Together, these coordinated transcriptional changes suggest that Hep_Bi-zonal cells undergo extensive metabolic remodeling that is compatible with enhanced stress adaptation, regenerative competence and increased cellular plasticity.

Because enrichment of peroxisome-associated programs emerged one of the most prominent features distinguishing tumor-fated hepatocytes, we next asked whether this transcriptional program is similarly engaged during human fatty liver disease progression. Across two independent human MASLD cohorts^130,131^, peroxisome-associated signature scores were calculated as described in Methods. Multiple peroxisome-associated signatures, including peroxisomal biogenesis, ROS buffering, lipid detoxification, and PPAR signaling, displayed their highest scores at the earliest steatotic stage (MAFL-F0) and remained consistently elevated throughout subsequent disease progression (Figure 5b; Supplementary Table 9). This pattern was recapitulated in an independent human Xenium dataset^132^, in which spatial signature scores were computed as described in Methods and revealed preferential enrichment of peroxisome-associated signatures along the midlobular-periportal axis, coinciding with regions of prominent lipid accumulation (Figure 5b; Supplementary Figure 8a). Notably, individual patient specimens displaying elevated peroxisomal activity simultaneously exhibited higher injury-repair responses, inflammatory signaling, and progressively increased RegHPC and NeoHPC signature scores (Figure 5b). These observations indicate that activation of peroxisome-associated metabolic programs accompanies the earliest emergence of regenerative progenitor-like states during human fatty liver disease progression.

Peroxisomes play central roles in very-long-chain fatty-acid β-oxidation and cellular redox homeostasis^128,129,133,134^.Recent work further demonstrated that peroxisomes directly protect mitochondrial function through stress-induced membrane contacts that mediate the transfer of mitochondrial reactive oxygen species into the peroxisomal lumen, thereby preserving mitochondrial redox homeostasis during oxidative stress^135^.Together with our observations in human disease, these findings raised the possibility that enhanced peroxisomal activity enables hepatocytes to adapt to the chronic lipotoxic, oxidative, and inflammatory stresses associated with fatty liver progression. If this model is correct, key components of the peroxisomal β-oxidation machinery would be expected to be selectively activated in tumor-fated hepatocytes. Consistent with this prediction, differential expression analysis identified Acox1, the first and rate-limiting enzyme of straight-chain very-long-chain fatty-acid β-oxidation in peroxisomes^98,136–138^, as one of the most prominently upregulated genes in Hep_Bi-zonal cells (Figure 5c; Supplementary Table 9). We therefore hypothesized that ACOX1-mediated peroxisomal metabolism represents a critical metabolic checkpoint governing hepatocyte survival and fate transition during liver tumor initiation.

To directly test this hypothesis, we engineered AKT-EGFP- and NRAS-tdTomato-expressing transposons to carry either non-targeting or Acox1-targeting sgRNAs, thereby genetically coupling oncogene expression with Acox1 perturbation in the same hepatocytes (Supplementary Figure 8b). Hepatocytes co-expressing EGFP and tdTomato were considered oncogene-expressing hepatocytes for all subsequent analyses. The constructs were introduced together with SB100X transposase by hydrodynamic tail-vein injection (Figure 5d; Supplementary Figure 8b; see Methods). Nineteen days after hydrodynamic transfection, mice receiving non-targeting sgRNAs developed extensive liver tumors, whereas livers from mice receiving Acox1 sgRNAs remained markedly smaller and grossly resembled those of healthy controls (Figure 5d). Consistent with these macroscopic observations, the liver-to-body weight ratio was significantly reduced following Acox1 disruption (0.178 ± 0.047 versus 0.298 ± 0.018 g/g in the non-targeting group, adjusted *P* < 0.0001), although it remained higher than that of healthy mice (0.050 ± 0.008 g/g, adjusted *P* < 0.0001) (Figure 5d; Supplementary Figure 8d). Flow cytometric analysis further confirmed a marked reduction in the proportion of oncogene-expressing hepatocytes following CRISPR-mediated disruption of Acox1, decreasing from an average of 16.0% in the non-targeting group to 2.57% in the Acox1 sgRNA group (Welch’s corrected two-tailed *t*-test, *P* < 0.0001; Figure 5e). Consistent with the substantial reduction in tumor burden, mice in the Acox1 sgRNA group showed significantly prolonged survival compared with those in the non-targeting sgRNA group. Five of seven mice in the non-targeting group reached the survival endpoint between Days 20 and 27, yielding a median survival of 24 days, whereas no deaths were observed among the seven Acox1-targeted mice during the corresponding follow-up period and median survival was not reached (log-rank Mantel-Cox test, χ² = 5.233, *P* = 0.0222; Figure 5f). Together, these findings demonstrate that ACOX1 is functionally required for efficient liver tumor development.

To determine when ACOX1 becomes essential during disease progression, we longitudinally monitored oncogene-expressing hepatocytes using HA immunohistochemistry together with immunofluorescence analysis (Figure 5g; Supplementary Figure 9a-d; see Methods). Comparable proportions of oncogene-expressing hepatocytes (EGFP⁺tdTomato⁺) were detected in the non-targeting and Acox1 sgRNA groups at Day 4 (27.3% versus 23.4%, adjusted *P* = 0.4507), indicating that Acox1 disruption did not affect the initial establishment of oncogene-expressing hepatocytes. However, whereas the proportion of oncogene-expressing hepatocytes progressively increased in control mice (32.8%, 38.5%, and 83.6% at Days 9, 14, and 19, respectively), it remained markedly reduced in Acox1-targeted livers (17.9%, 16.4%, and 7.1%, respectively), with significant differences emerging at Day 9 (adjusted *P* < 0.0001), Day 14 (adjusted *P* < 0.0001), and Day 19 (adjusted *P* < 0.0001). These findings indicate that ACOX1 is dispensable for the initial establishment of oncogene-expressing hepatocytes but becomes essential for their subsequent survival and clonal expansion during fatty liver-associated liver tumor initiation.

To further validate this requirement independently of CRISPR-mediated gene editing, we inhibited ACOX1 using the reported small-molecule inhibitor 10,12-tricosadiynoic acid (TDYA) ^139^ and established an oncogene-induced conditional Acox1 deletion model (Acox1^ΔHep) by hydrodynamically delivering Cre-expressing AKT and NRAS vectors into Acox1^fl/fl mice (Supplementary Figure 8f-g; see Methods). Consistent with the CRISPR-mediated perturbation, both pharmacological inhibition and conditional genetic deletion of Acox1 markedly reduced the abundance of oncogene-expressing hepatocytes (Supplementary Figure 8e), demonstrating that the requirement for ACOX1 is reproducible across independent experimental strategies. Together, these complementary genetic and pharmacological approaches demonstrate that ACOX1-dependent peroxisomal metabolism is required for efficient liver tumor initiation.

To investigate the molecular consequences of ACOX1 disruption, we longitudinally profiled the transcriptomes of FACS-isolated oncogene-expressing hepatocytes (EGFP⁺tdTomato⁺) from the CRISPR-sgRNA model at Days 4, 9 and 14, comparing cells carrying non-targeting or Acox1-targeting sgRNAs (Figure 5h-j; Supplementary Figure 8b, d; Supplementary Table 9; see Methods). Signature analysis revealed that oncogene-expressing hepatocytes carrying Acox1-targeting sgRNAs largely retained mature hepatocyte transcriptional programs, whereas oncogene-expressing hepatocytes carrying non-targeting sgRNAs progressively acquired Hep_Bi-zonal and RegHPC signatures (Figure 5h). Consistently, principal component analysis showed that Acox1-targeted hepatocytes remained transcriptionally closer to normal hepatocytes throughout disease progression, whereas control cells progressively diverged toward a distinct tumor-associated trajectory (Figure 5i). Longitudinal GSEA further demonstrated that Acox1 disruption markedly attenuated the transcriptional remodeling associated with the tumor-fated lineage (Figure 5j). Compared with control cells, Acox1-targeted hepatocytes showed reduced activation of antioxidant defense, inflammatory signaling, regenerative and progenitor-state programs, while retaining higher activities of mature hepatic metabolic pathways together with persistent mitochondrial dysfunction, oxidative stress and apoptotic signatures. Collectively, these findings identify ACOX1-dependent peroxisomal β-oxidation as a previously unrecognized metabolic checkpoint required for the coordinated transcriptional and metabolic remodeling that enables hepatocyte adaptation, progenitor-state formation, and subsequent tumor initiation.

### Spatial remodeling identifies a macrophage niche associated with tumor-fated hepatocytes

Having identified ACOX1-dependent peroxisomal metabolism as a critical intrinsic mechanism enabling hepatocytes to adapt to the metabolic stresses associated with fatty liver progression, we next asked whether extrinsic cues from the surrounding tissue microenvironment further influence hepatocyte fate decisions during tumor initiation. Because liver zonation establishes a highly organized metabolic landscape and immune cells actively participate in tissue injury and regeneration^24,26,140^, we hypothesized that spatial organization and local cellular interactions cooperatively shape the emergence of tumor-fated hepatocyte states. To address this hypothesis, we performed longitudinal spatial transcriptomic profiling across four stages of liver tumor initiation (Day 4, Day 9, Day 14, and Day 19) (Figure 6; Supplementary Table 10; see Methods).

**Figure 6.**
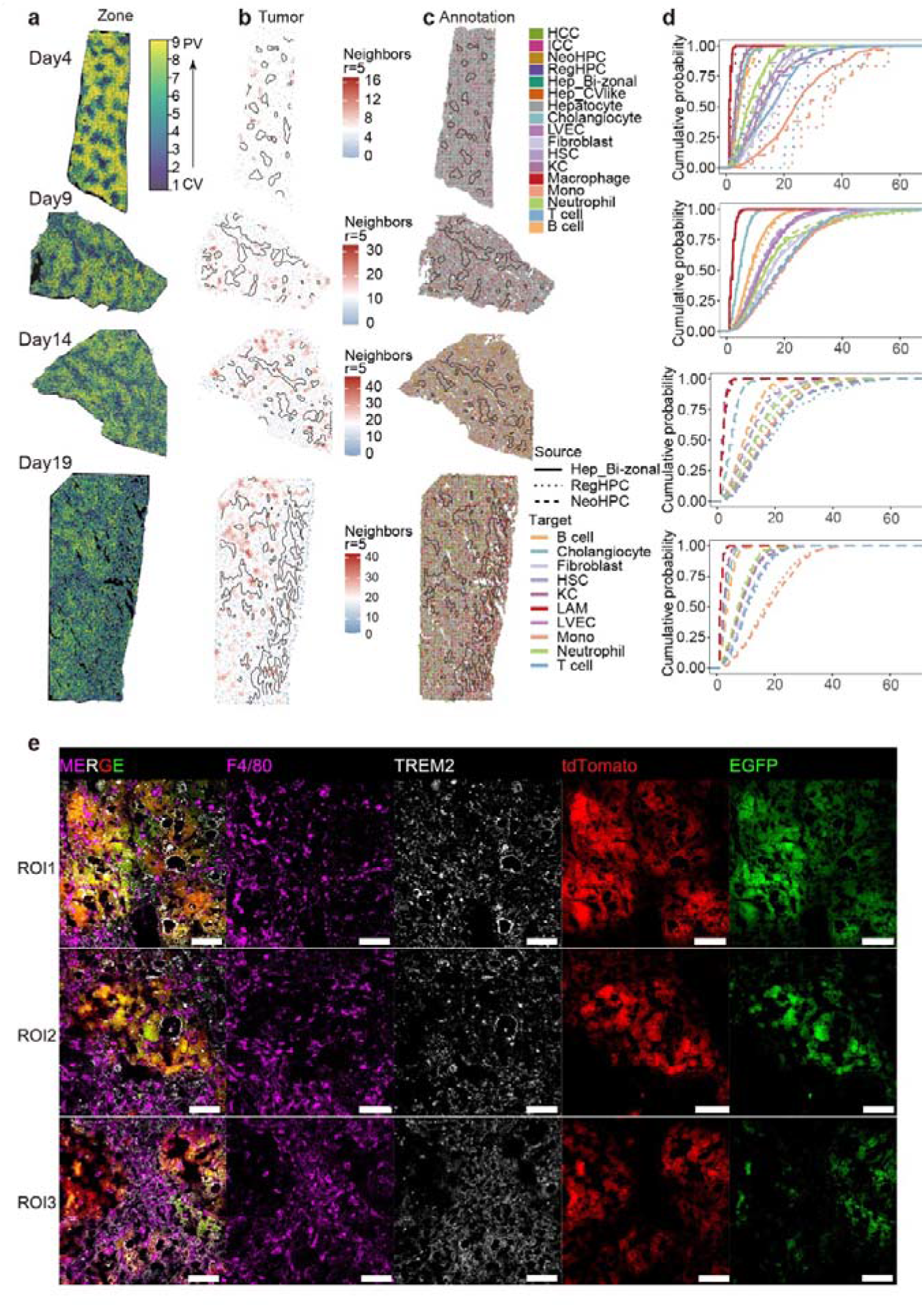
Spatial transcriptomics identifies macrophage-associated niches around tumor-fated hepatocytes. **(a)** Spatial maps showing liver zonation coordinates at Days 4, 9, 14, and 19 after induction. The tissue sections are divided into nine continuous layers spanning the pericentral (CV, layer 1, purple) to periportal (PV, layer 9, yellow) regions. **(b)** Spatial density maps of oncogene-expressing cells at Days 4, 9, 14, and 19. Local cell density was calculated using a neighborhood search radius of five nearest neighbors and is shown from low (blue) to high (red). Black outlines indicate the pericentral (CV) regions. **(c)** Spatial cell-state annotation maps corresponding to the tissue sections shown in (a) and (b). Cells are colored according to their annotated identities, including Hep_CVlike, Hep_Bi-zonal, RegHPC, NeoHPC, HCC, ICC, Kupffer cells (KC), macrophages, neutrophils, endothelial cells (LVEC), fibroblasts, and other annotated cell populations. Black outlines indicate the pericentral (CV) regions. **(d)** Nearest-neighbor distance analysis between hepatocyte states and microenvironmental cell populations. Curves show the cumulative probability distribution as a function of the distance between Hep_Bi-zonal (solid line), RegHPC (dotted line), or NeoHPC (dashed line) cells and the indicated stromal or immune cell populations. Colors denote the indicated target cell types. **(e)** Representative multiplex immunofluorescence images from three independent regions of interest (ROI1-ROI3). Separate and merged channels are shown for F4/80 (magenta), TREM2 (white), tdTomato (red), EGFP (green), and DAPI (blue). Scale bars, 50 μm.

Using the single-cell reference atlas, we annotated cell identities across spatial sections and observed high concordance between the transcriptional profiles of hepatocyte populations identified by single-cell RNA sequencing and their corresponding spatially mapped counterparts, supporting robust integration of the two datasets (Figure 6c; Supplementary Figure 10b, c).

We next reconstructed the hepatic zonation axis to determine how normal liver architecture changes during tumor initiation. Spatial spots were assigned to nine reconstructed lobular layers using a landmark gene-based computational framework^72^(see Methods). At Day 4, the normal lobular architecture remained remarkably well preserved, exhibiting sharply defined zonation boundaries and a well-organized metabolic zonation pattern across the hepatic lobule (Figure 6a; Supplementary Figure 10a-c). As disease progressed from Day 9 to Day 19, however, the boundaries separating the pericentral, midlobular, and periportal compartments became progressively disrupted and were eventually largely lost, accompanied by a gradual decline in the expression of representative zonation markers, including Cyp2e1, Hamp2, and Gls2 (Figure 6a; Supplementary Figure 10a-c). These observations indicate that progressive remodeling of the hepatic zonation architecture accompanies fatty liver-associated liver tumor initiation.

We then examined where tumor-fated hepatocyte states emerge within the liver lobule. Spatial neighborhood analysis revealed that tumor hotspots were consistently enriched outside the pericentral region and preferentially localized along the midlobular-periportal axis (Figure 6b). Spatial annotation based on the single-cell reference further resolved Hep_Bi-zonal, RegHPC, NeoHPC, HCC, and other hepatocyte states within these tissue sections, enabling subsequent analyses of their spatial organization and interactions with the surrounding microenvironment (Figure 6c). These spatial observations independently corroborate the lineage-tracing results presented in Figure 3, providing orthogonal evidence that Hep_Bi-zonal, rather than Hep_CVlike, represents the principal tumor-fated hepatocyte population. Consistently, multiplex immunofluorescence demonstrated that expanding EGFP tdTomato lesions were largely excluded from GLUL pericentral regions, further confirming that tumor initiation preferentially occurs outside the central vein niche (Supplementary Figures 9a-d and 10f).

Because inflammatory signaling and myeloid interaction programs were prominent features of RegHPC, we investigated whether immune cells were spatially associated with tumor-fated hepatocyte populations during disease progression (Figure6d, e). Nearest-neighbor analysis^141^ revealed that lipid-associated macrophages (LAMs) consistently represented the nearest immune population to Hep_Bi-zonal, RegHPC, and NeoHPC cells across multiple stages of tumor initiation (Figure 6d; Supplementary Table 10; See Methods). Moreover, the spatial proximity between LAMs and these transitional hepatocyte populations progressively increased as disease advanced, suggesting the gradual formation of a macrophage-enriched spatial niche surrounding tumor-fated hepatocytes. Multiplex immunofluorescence independently confirmed this spatial organization, demonstrating prominent accumulation of TREM2 F4/80 macrophages immediately adjacent to oncogene-expressing hepatocytes (EGFP^+^tdTomato^+^) across multiple independent regions of interest (Figure 6e).

Collectively, these findings demonstrate that fatty liver-associated tumor initiation is accompanied by progressive disruption of the normal hepatic zonation architecture, within which tumor-fated hepatocytes emerge preferentially along the midlobular-periportal axis and progressively establish a spatially restricted macrophage-enriched niche.

### Dual ontogenies and functional specialization of lipid-associated macrophages shape the tumor-fated hepatocyte niche

Having observed that lipid-associated macrophages (LAMs) were spatially enriched around tumor-fated hepatocyte populations, we next sought to define their developmental origins, temporal dynamics, and functional contributions during liver tumor initiation. The hepatic myeloid compartment comprises tissue-resident Kupffer cells (KCs), infiltrating monocytes, and monocyte-derived or differentiated macrophage populations, which collectively participate in lipid handling, inflammatory responses, tissue injury, and repair.^65,67,142,143^ High-resolution subclustering of myeloid cells resolved three major compartments including KCs, monocytes, and macrophages, with distinct transcriptional identities confirmed by signature scoring and canonical marker expression^65,144^ (Figure 7a; Supplementary Figure 11a-c; Supplementary Table 11).

**Figure 7.**
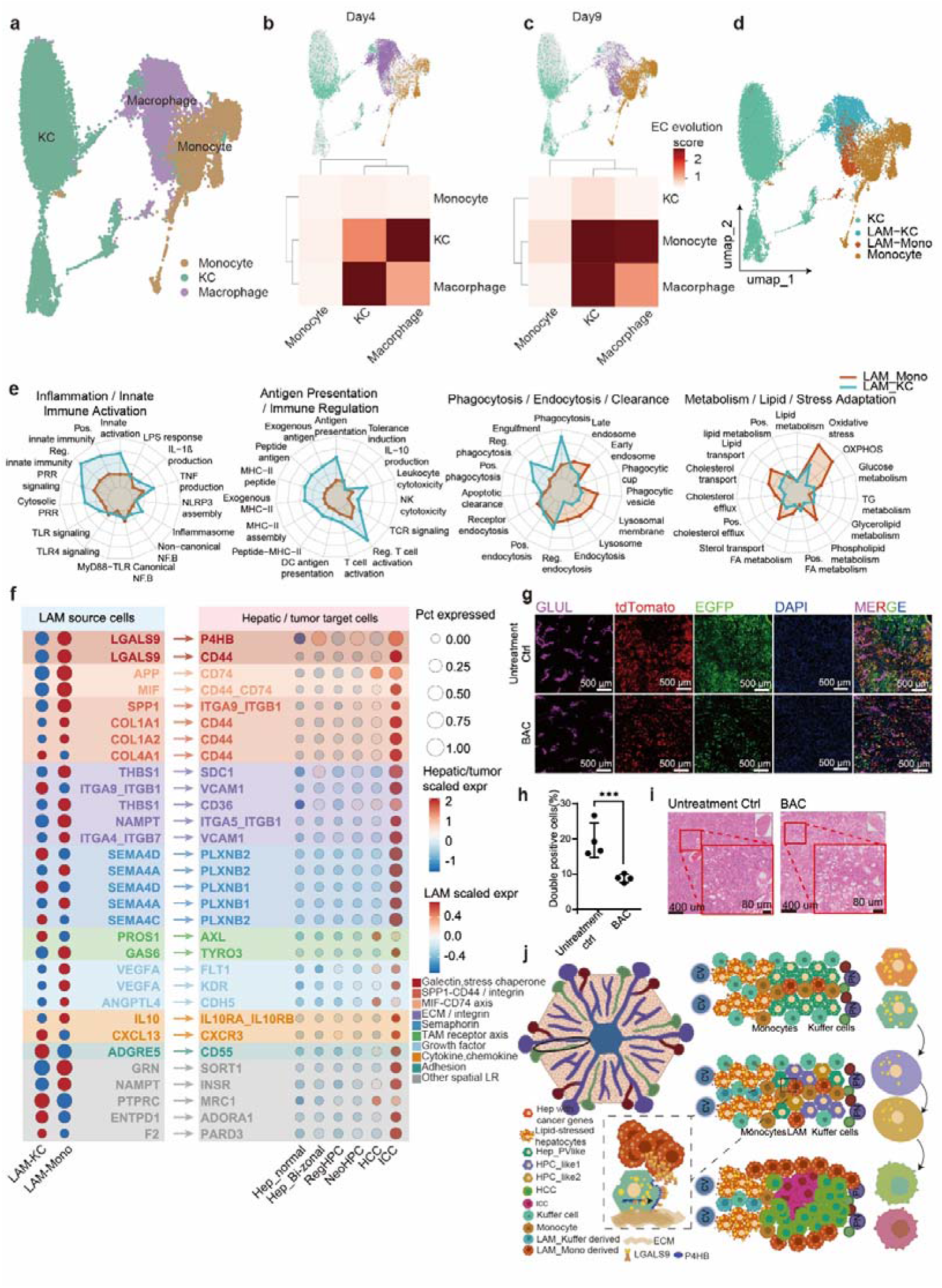
Dual ontogenies and functional specialization of lipid-associated macrophages during liver tumor initiation. **(a)** UMAP visualization of hepatic myeloid cell populations, including Kupffer cells (KCs), monocytes, and macrophages. **(b, c)** Heatmaps showing normalized evolutionary coupling (EC) scores among myeloid cell populations at Day 4 (b) and Day 9 (c). Colors indicate normalized EC scores. **(d)** UMAP visualization of lipid-associated macrophage (LAM) populations, including Kupffer cell-derived LAMs (LAM-KC), monocyte-derived LAMs (LAM-Mono), and their corresponding precursor populations. **(e)** Radar plots showing representative enriched GO and KEGG pathways for LAM-KC and LAM-Mono cells. Enriched pathways are grouped into inflammation/innate immune response, antigen presentation/immune regulation, phagocytosis/endocytosis, and metabolism/lipid-related categories. **(f)** Dot plot showing ligand-receptor interactions between LAM populations (LAM-KC and LAM-Mono) and hepatocyte states, including Hep_Normal, Hep_Bi-zonal, RegHPC, NeoHPC, HCC, and ICC. Ligand-receptor pairs are grouped into the indicated signaling categories, including galectin, SPP1-CD44/integrin, MIF-CD74, extracellular matrix (ECM)/integrin, semaphorin, TAM receptor, growth factor, cytokine/chemokine, and adhesion pathways. Dot size indicates the percentage of cells expressing the indicated ligand or receptor, and colors indicate scaled expression levels in sender and receiver cell populations. **(g)** Representative multiplex immunofluorescence images of liver sections from untreated control and bacitracin (BAC)-treated mice. Separate and merged channels are shown for GLUL (magenta), tdTomato (red), EGFP (green), and DAPI (blue). Scale bars, 500 μm. The experimental design is shown in Supplementary Figure 8h. **(h)** Quantification of EGFP⁺tdTomato⁺ cells corresponding to the representative images shown in (g). Data were analyzed using a two-tailed unpaired *t* test with Welch’s correction (n = 4 mice per group; *P* = 0.0215). **(i)** Representative hematoxylin and eosin (H&E) staining of liver sections from untreated control and BAC-treated mice. Scale bars, 400 μm (main images) and 80 μm (enlarged views). **(j)** Schematic illustrating macrophage populations, hepatocyte states, and the indicated ligand-receptor interactions across the liver lobule. PN = portal node (a vasculature structure composed from the portal vein (purple), hepatic artery (red) and bile duct (green)), CV = central vein (blue).

To infer the developmental relationships among these myeloid populations, we calculated evolutionary coupling scores across early stages of tumor initiation (Figure 7b, c). At Day 4, newly emerging macrophages displayed markedly stronger evolutionary coupling with resident KCs than with monocytes, suggesting that the earliest macrophage response was primarily associated with the tissue-resident Kupffer-cell compartment. By contrast, at Day 9, macrophages showed substantially increased evolutionary coupling with monocytes, indicating that the sustained macrophage response during later tumor initiation was progressively supplied by recruited circulating monocytes (Figure 7b, c). Based on these inferred lineage relationships and their transcriptional signatures, we separated the LAM compartment into two ontogenetically distinct subsets: KC-associated LAMs (LAM-KC) and monocyte-associated LAMs (LAM-Mono) (Figure 7d; Supplementary Figure 11c).

We further validated these computationally inferred origins using an independent Ms4a3-Cre; Rosa26-LSL-tdTomato genetic lineage-tracing model, which enables permanent labeling of monocyte-lineage cells and their progeny^145^. (Supplementary Figure 11d; see Methods). To specifically analyze intrahepatic immune cells, livers were carefully perfused before tissue collection to minimize contamination by circulating intravascular immune cells. In healthy control livers, monocyte-derived macrophages represented only a minor fraction of the hepatic myeloid compartment. Following oncogene induction, however, the hepatic macrophage compartment underwent a striking compositional shift. Monocyte-derived macrophages, which accounted for only 2.6% of hepatic macrophages in healthy liver, expanded rapidly to approximately 30% by Day 4 and reached parity with resident Kupffer cells by Day 7 (50.1% versus 49.9%) (Supplementary Figure 11e). By Day 14, monocyte-derived macrophages had become the predominant macrophage population (78.8%), remaining the major component at Day 21 (60.6%) (Supplementary Figure 11e). A similar transition was observed within the TREM2 LAM compartment (Supplementary Figure 11f). Whereas early LAMs were primarily KC-derived (76.3% LAM-KC) under homeostatic conditions and remained largely KC-associated at Day 4 (65.9%), monocyte-derived LAMs progressively expanded thereafter, exceeding KC-derived LAMs by Day 7 (63.3% versus 36.7%) and ultimately constituting nearly 90% of all LAMs by Day 21 (88.7%). These findings independently validate the evolutionary coupling analysis and further demonstrate that the LAM compartment undergoes a sequential ontogenetic transition from resident KC-derived macrophages to monocyte-derived macrophages during liver tumor initiation.

We next asked whether LAM-KC and LAM-Mono populations exhibited distinct functional programs. Functional profiling revealed a clear division of labor between LAM-KC and LAM-Mono subsets (Figure 7e; Supplementary Table 11). LAM-KC cells were preferentially enriched for innate immune activation, Toll-like receptor signaling, inflammatory responses, antigen presentation, and phagocytic programs, consistent with the established sentinel and debris-clearance functions of tissue-resident Kupffer cells^67,143^. In contrast, LAM-Mono cells were enriched for endocytosis, lysosomal activity, lipid uptake, cholesterol transport, and stress-adaptation programs, suggesting specialization in lipid handling, clearance of damaged material, and remodeling of the evolving tumor-associated niche^67,69,146,147^. Spatially resolved pathway analysis further supported this temporal specialization: early macrophage-enriched regions were characterized by complement activation, autophagy-associated remodeling, and tissue-migration programs, whereas later regions showed stronger enrichment of extracellular matrix organization, leukocyte chemotaxis, antigen presentation, lipid metabolism, and phagocytosis-associated pathways (Supplementary Figure 11h). Together, these data suggest that LAM-KC cells may function as early inflammatory and phagocytic responders, whereas LAM-Mono cells progressively establish a lipid-handling and remodeling niche that associated with tumor-fated hepatocyte expansion.

Given the strong spatial association between LAMs and tumor-fated hepatocytes observed in Figure 6d, we next examined whether the two LAM subsets differed in their spatial relationships with hepatocyte-derived transitional states. Nearest-neighbor analysis revealed that Hep_Bi-zonal, RegHPC, and NeoHPC cells were consistently positioned in closer proximity to LAM-Mono than to LAM-KC cells, particularly during the later stages of tumor initiation (Supplementary Figure 11g). This preferential spatial association prompted us to investigate whether monocyte-derived LAMs establish distinct intercellular communication with tumor-fated hepatocytes.

To identify candidate signaling mechanisms underlying this spatial association, we integrated spatial ligand-receptor activity inferred using LIANA^148,149^ with communication analysis in an aggregated single-cell transcriptomic reference, as described in Methods. This analysis identified a prominent signaling interaction between LAM-Mono cells and tumor-fated hepatocyte states through the LGALS9-P4HB signaling axis (Figure 7f; Supplementary Table 11; see Methods). LGALS9 was preferentially expressed by LAM populations, particularly LAM-Mono cells, whereas P4HB was broadly expressed by hepatocyte-derived states, including Hep_Bi-zonal, RegHPC, NeoHPC, HCC, and ICC. P4HB encodes protein disulfide isomerase A1 (PDIA1), an endoplasmic-reticulum-resident oxidoreductase and molecular chaperone that catalyzes disulfide bond formation during protein folding ^150,151^. Previous studies have shown that, under conditions of cellular stress, a fraction of P4HB can translocate to the plasma membrane, where it interacts with extracellular LGALS9 and contributes to the stabilization of membrane-associated P4HB^151–154^. These observations identified the LGALS9-P4HB axis as a candidate ligand-receptor signaling axis linking monocyte-derived LAMs with tumor-fated hepatocyte populations. To determine whether P4HB contributes functionally to tumor initiation, we treated mice with bacitracin (BAC), a pharmacological inhibitor of P4HB^156,157^. BAC treatment markedly reduced the abundance of oncogene-expressing hepatocytes, with EGFP⁺tdTomato⁺ cells decreasing from an average of 19.6% in untreated mice to 9.3% following BAC treatment (Welch’s corrected two-tailed *t*-test, *P* = 0.0215; Figure 7g-i, Supplementary Figure 8h). Histological analyses further confirmed a pronounced reduction in both premalignant and malignant lesion formation after P4HB inhibition, consistent with the marked suppression of tumor initiation observed by immunofluorescence (Figure 7i; Supplementary Figure 8e). Together, these findings demonstrate that P4HB activity is required for efficient expansion of tumor-fated hepatocytes during liver tumor initiation.

Together, these observations indicate that progressive replacement of resident KC-derived LAMs by monocyte-derived LAMs is accompanied by functional specialization and establishment of the LGALS9-P4HB signaling axis, thereby creating a distinct macrophage-associated niche that supports tumor-fated hepatocyte expansion during liver tumor initiation (Figure 7j).

## Discussion

This study defines a lineage-resolved conceptual framework governing fatty liver-associated hepatocarcinogenesis. By coupling CERAMIC lineage recording with longitudinal single-cell and spatial transcriptomics, we distinguish transiently altered, reactive hepatocyte states from those that are genuinely tumor-fated. Resolving this divergence is fundamental to identifying the true cells of origin in liver cancer. Disrupted or chronically injured tissues frequently harbor a broad reservoir of abnormal, progenitor-like, or stress-adapted cells; however, only a subset of these cells ultimately contributes to tissue regeneration or malignant clonal expansion^21,28,30,72,93,99^. Our data establish Hep_Bi-zonal cells as the pivotal gateway to a tumor-fated hierarchy that proceeds sequentially through regenerative and neoplastic hepatocyte progenitor states, ultimately diversifying into both HCC and ICC lineages. In doing so, this study moves beyond a static cataloging of premalignant states to reconstruct how cell state, lineage history, spatial coordinates, and microenvironmental niche support are dynamically coupled during the earliest phases of tumor initiation.

The identification of the Hep_Bi-zonal state as a major tumor-fated population introduces a critical dimension to emerging models of zone-dependent liver tumorigenesis. While recent work indicates that baseline hepatocyte zonation biases transformation potential^35,38,39^, metabolic zonation alone cannot fully explain why certain oncogene-expressing hepatocytes expand clonally while neighboring, identically driven cells remain non-progressive or fail to transform. In our model, Hep_Bi-zonal and Hep_CVlike cells represent two divergent responses to the same oncogenic and metabolic context. Although both populations experience chronic lipotoxic and inflammatory stress, Hep_CVlike cells largely fail to progress beyond a stress-adapted state, whereas Hep_Bi-zonal cells exhibit continuous lineage progression through RegHPC and NeoHPC intermediates toward malignant descendants. This distinction extends recent observations that experimentally induced plastic hepatocytes can adopt tumor-restrictive states that are refractory to proliferative and oncogenic cues^28^. Rather than suggesting that hepatocyte plasticity is uniformly protective or uniformly tumor-promoting, our lineage-resolved analyses indicate that plasticity itself is heterogeneous. Specifically, only a metabolically adaptive Hep_Bi-zonal state acquires sustained progenitor-like plasticity and ultimately seeds the tumor-fated hierarchy, whereas alternative stress-associated states remain lineage-restricted. These findings suggest that transformation competence is not determined solely by lobular position or injury-induced plasticity, but instead emerges from the coordinated integration of spatial identity, metabolic adaptation, and progressive lineage reprogramming.

A key implication of this cellular hierarchy is that metabolic adaptation acts as a critical gatekeeper for fate progression. Metabolic dysfunction-associated steatohepatitis exposes hepatocytes to chronic lipotoxic and oxidative stress^10,11,158^, creating selective pressures that eliminate damaged cells, induce senescence, or favor the persistence of stress-resistant populations. Our data identify ACOX1-dependent peroxisomal-oxidation as an absolute requirement for the survival and subsequent progenitor-state transition of tumor-fated hepatocytes. This finding extends the classical view of peroxisomes in lipid homeostasis and redox regulation into the earliest, morphologically cryptic phases of liver initiation. Rather than functioning merely as a constitutive lipid-processing pathway, peroxisomal metabolism appears to sustain a highly selective stress-adaptation program that supports persistence of Hep_Bi-zonal cells while enabling progressive lineage reprogramming. Although the precise biochemical route remains to be fully dissected^127,129,159^, such as whether ACOX1 primarily supports tumor-fated cells by detoxifying specific lipotoxic species, modulating localized reactive oxygen species, reshaping cellular redox balance, altering lipid-droplet flux, or coordinating a combination of these processes, our perturbation data establish a clear functional dependence. Future work should uncouple these candidate mechanisms and determine whether an identical peroxisomal metabolic checkpoint operates in human MASLD/MASH-associated premalignant hepatocytes.

The spatial architecture of tumor-fated hepatocytes further implies that the liver lobule functions as a topographical selection landscape during tumor initiation. In homeostasis, hepatocyte functions are distributed along the porto-central axis^24,26^, forming a highly organized metabolic zonation that undergoes profound remodeling during injury and carcinogenesis^43,72,93^. Our spatial analyses reveal that tumor-fated hepatocyte states preferentially expand along the midlobular-to-periportal axis, becoming progressively incorporated within macrophage-rich microenvironments as initiation advances. This non-random pattern is unlikely to represent a passive reflection of expanding tumor burden. Instead, it strongly indicates that specific tissue territories provide a distinct combination of metabolic, inflammatory, and regenerative signals that actively support clonal persistence. Concurrently, the direction of causality warrants cautious interpretation. Pre-existing spatial positioning may actively shape the induction of the Hep_Bi-zonal state, or alternatively, tumor-fated Hep_Bi-zonal cells may selectively expand into territories that are subsequently remodeled around them. Distinguishing between these dual possibilities will necessitate temporal perturbations that alter local zonation, injury gradients, or niche composition prior to definitive lineage commitment.

The macrophage data delineate a second, cell-extrinsic interaction layer that structurally and molecularly couples with the hepatocyte compartment. Although lipid-associated macrophages (LAMs) have been documented across various contexts of metabolic disease, tissue repair, and established cancer^22,66,67,77,147^, the collective designation can obscure critical differences in cellular origin, recruitment temporal dynamics, and specialized functions. Our longitudinal analysis successfully separates an early, resident Kupffer cell (KC)-associated LAM response from a late-acting, bone marrow monocyte-derived LAM population that preferentially exhibits a strong spatial co-localization with tumor-fated hepatocyte states. This temporal replacement demonstrates that macrophage remodeling during tumor initiation is not a uniform inflammatory response. KC-derived LAMs may primarily participate in initial injury sensing, phagocytosis, and tissue clearance, whereas monocyte-derived LAMs are preferentially enriched for pathways related to lipid handling, lysosomal activity, and molecular engagement within the evolving pre-neoplastic niche. This intra-lineage division of labor aligns with emerging paradigms showing that recruited and resident hepatic macrophages can adopt overlapping LAM-like phenotypes while executing divergent functional programs in tissue injury^67^. Crucially, in the tumor-initiation setting, this late-recruited monocyte-derived LAM state becomes preferentially coupled to hepatocyte fate progression in spatial and transcriptomic coordinates rather than simply tracking generic damage resolution.

Mechanistically, the LGALS9-P4HB signaling axis emerges as a prominent candidate pathway through which monocyte-derived LAMs support tumor-fated hepatocytes. The prominent expression of LGALS9 in LAM populations, the upregulation of P4HB across tumor-fated hepatocyte states, the localized spatial ligand-receptor activity, and the notable suppression of tumor expansion via pharmacological P4HB inhibition collectively provide convergent evidence for a functional link between this microenvironmental signal and early tumor growth. Nonetheless, this interpretation should remain appropriately cautious. While current evidence demonstrates that P4HB activity is required for efficient clonal expansion in this model and highlights LGALS9-P4HB as a key interaction candidate connecting LAM-Mono cells to tumor-fated hepatocytes, direct ligand-receptor causality requires further genetic dissection. Definitive proof will necessitate cell-type-specific perturbations, including selective genetic deletion of *Lgals9* in LAM-Mono cells combined with hepatocyte-specific manipulation of *P4hb*, to test whether disrupting this physical axis directly compromises hepatocyte stress resistance, progenitor-state maintenance, or clonal expansion. Even within these boundaries, our findings conceptually bridge immune-niche remodeling with the cell-autonomous metabolic adaptations that enable tumor-fated hepatocytes to survive and expand during the earliest stages of liver tumor initiation.

Collectively, these insights converge on a cooperative model wherein fatty liver-associated tumor initiation is driven by the intersection of hepatocyte-intrinsic and microenvironmental programs. Oncogenic activation alone is insufficient to account for the extraordinary selectivity of malignant transformation. Rather, tumor-fated hepatocytes coordinately acquire a permissive transcriptional state, withstand severe metabolic injury via peroxisomal adaptation, expand within a supportive lobular territory, and receive instructive niche signals from ontogenetically remodeled macrophages. This cooperative model effectively reconciles two traditionally views of early liver cancer, where one is centered on cell-autonomous metabolic and genetic rewiring and the other is focused on inflammatory or stromal remodeling. Our data indicate that these internal metabolic shifts and external immune responses act in concert remarkably early, intertwining long before tumors become visible to dictate which hepatocytes ultimately acquire tumor fate.

Several limitations should be considered. While the AKT/NRAS model^81^ faithfully recapitulates core clinical features of fatty liver-associated mixed liver tumor initiation, it does not capture all human etiologies, alternative oncogenic drivers, or the protracted timescales characteristic of human HCC and ICC development. Accordingly, the generalizability of the Hep_Bi-zonal trajectory, the dependency on ACOX1, and the supportive role of the LAM-Mono niche must be rigorously evaluated across additional models of MASLD/MASH, fibrosis-driven carcinogenesis, and human premalignant tissue specimens. Furthermore, while our lineage evidence points to a shared upstream tumor-fated trajectory that ultimately diversifies into both HCC and ICC lineages, the precise molecular branching rules as well as cell-intrinsic or cell-extrinsic cues that govern the final HCC-versus-ICC fate choice require deeper investigation. Finally, although small-molecule P4HB inhibition underscores the functional relevance of the niche, it does not establish the cell-type-specific contribution of P4HB or prove an absolute requirement for the upstream LGALS9 ligand.

Despite these limitations, this study provides a robust conceptual framework for therapeutic interception before established malignant lesions. These findings suggest that effective early prevention may require a paradigm shift, from targeting isolated oncogenic pathways or generalized inflammation to disrupting the metabolic adaptations and niche interactions that enable rare premalignant hepatocytes to acquire tumor fate. Peroxisomal

-oxidation checkpoints and macrophage-hepatocyte cross-talk pathways thus emerge as highly promising candidate therapeutic vulnerabilities for future prevention-oriented clinical studies. More broadly, the CERAMIC framework offers a generalizable strategy for resolving how tissue injury, spatial topology, and immune remodeling collectively shape the fate of rare premalignant cells. Ultimately, integrating lineage recording with spatial and molecular profiling provides a general framework for distinguishing transient injury responses from the rare cellular trajectories that ultimately give rise to cancer.

## Methods

Detailed experimental procedures and analytical methods will be made publicly available upon publication of the peer-reviewed article.

## Data availability

The sequencing datasets generated during this study are being deposited in the Gene Expression Omnibus (GEO) and will be publicly available upon publication. The GEO accession number will be included once available. All other data supporting the findings of this study are available from the corresponding authors upon reasonable request.

## Code availability

The custom scripts used for data processing and analysis are available from the corresponding author upon reasonable request.

## Acknowledgements

We thank Prof. Luo-Nan Chen for valuable advice on computational analyses and data interpretation. We thank Prof. Xuxu Sun (Shanghai Jiao Tong University) for generously providing the oncogenic plasmids and for valuable scientific discussions. We thank Prof. Guangchuan Wang (Center for Excellence in Molecular Cell Science, Chinese Academy of Sciences) for generously providing the Ms4a3-Cre mice. We thank Prof. Pengfei Sui (Center for Excellence in Molecular Cell Science, Chinese Academy of Sciences) for providing the Rosa26-LSL-tdTomato mice. We also thank all members of the Liu laboratory for helpful discussions and technical support.

## Author contributions

J.F. conceived the study, designed and performed experiments, analyzed and interpreted the data, and wrote the manuscript. P.J. performed data quality control, preprocessing, and computational analyses. X.N. assisted with mouse model generation, mouse genotyping, and bulk genomic DNA library preparation. S.M. performed the majority of mouse genotyping. X.W. managed reagent procurement and experimental logistics. L.C. supervised computational analyses, interpreted the data, and revised the manuscript. X.S. provided the oncogenic plasmids, contributed to study design, and provided scientific guidance. X.L. conceived and supervised the study, interpreted the data, secured funding, and revised the manuscript. All authors discussed the results and approved the final manuscript.

---

**Supplementary Figure 1.**
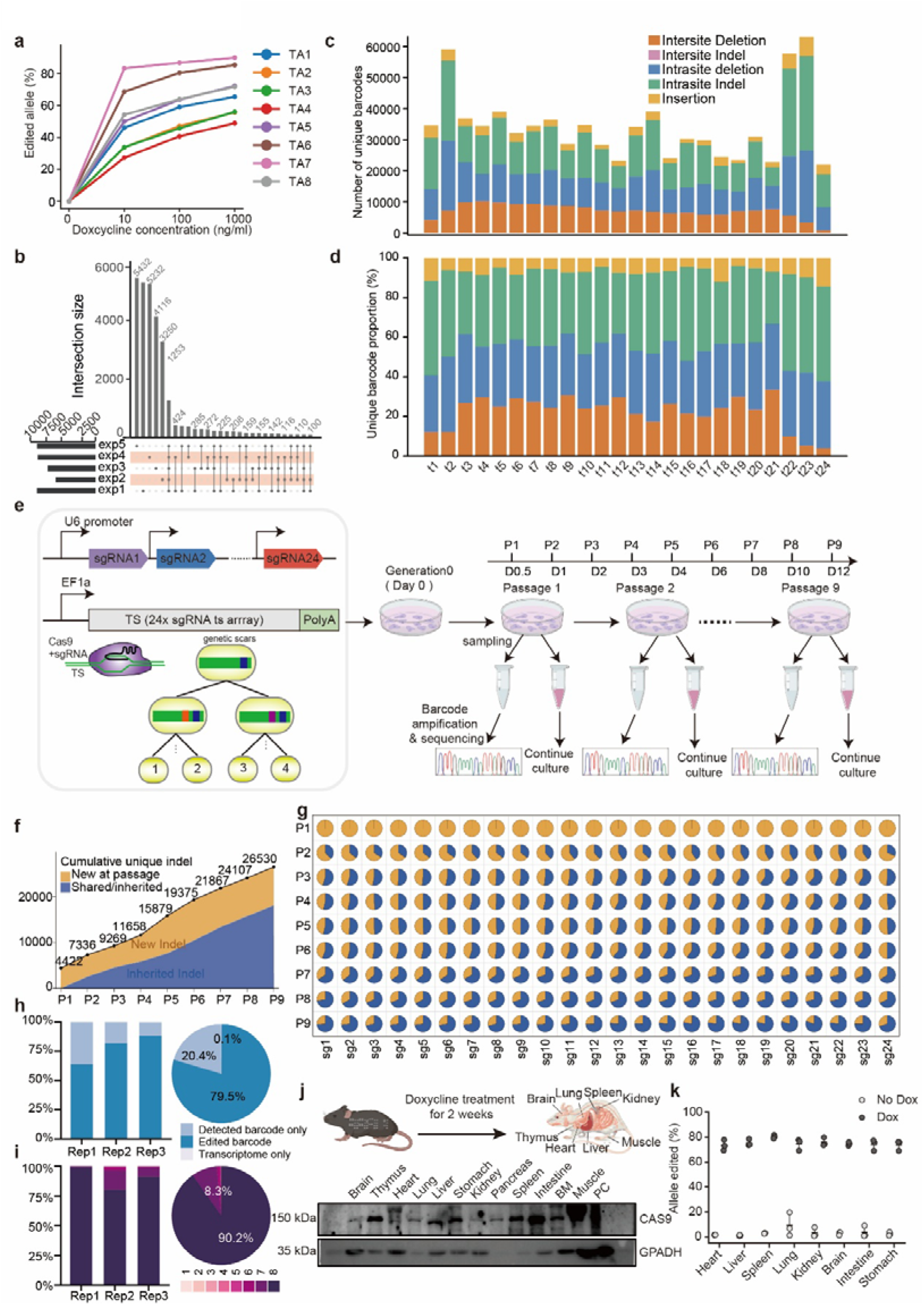
Technical validation of CERAMIC recording capacity and recovery. **(a)** Editing efficiency across the eight individual target array modules (TA1-TA8) responsive to doxycycline induction. The curves plot the editing frequencies calculated from CERAMIC mouse embryonic stem cells (ESCs) following 48 h of induction across a gradient of doxycycline concentrations. The y-axis represents the percentage of edited alleles (%), and the x-axis denotes the doxycycline doses (0, 10, 100, and 1000 ng/mL). Distinct color-coded lines represent the editing efficiencies of the eight target-array (TA) modules across the indicated doxycycline concentrations. **(b)** UpSet plot illustrating the intersection and overlap of unique edited barcode species across five independent experimental replicates (exp1-exp5) in CERAMIC mouse embryonic stem cells (ESCs) following 96 h of induction with 100 ng/mL doxycycline. The horizontal bars on the bottom-left denote the total barcode pool size captured per experiment; the vertical bars and upper labels indicate the exact intersection size for each overlapping matrix combination. **(c)** Absolute numbers of unique edited barcodes across target positions t1-t24. Barcodes containing deletions larger than 400 nt were excluded. The plot represents a comprehensive compilation of all edited patterns pooled from the ESC characterization assays (Supplementary Fig. S1a, e) and the in vivo single-cell and spatial transcriptomics datasets (Fig. 1d). Barcodes are color-coded according to five structural categories: intersite deletion (orange; >150 nt), intersite indel (purple; >150 nt deletion with concurrent insertions), intrasite deletion (blue; ≤150 nt), intrasite indel (green; concurrent insertions and ≤150 nt deletions), and insertion (yellow; insertion-only). **(d)** Relative percentage composition of the four unique edited barcode categories across target positions t1-t24, utilizing the identical aggregated multi-experiment dataset as in (c). The stacked bar chart shows the relative proportions of the four barcode categories across target positions t1-t24. **(e)** Experimental workflow of the serial-passaging assay using CERAMIC mouse embryonic stem cells (ESCs). The schematic on the left illustrates the multi-promoter array driving 24 sgRNAs paired with an EF1a-driven target array (TS), where continuous Cas9-mediated editing introduces genetic scars into the target array. At Generation 0 (Day 0), Cas9 expression is induced. Cells are sequentially passaged from P1 through P9 at defined intervals (D0.5, D1, D2, D3, D4, D6, D8, D10, and D12). **(f)** Cumulative numbers and composition of unique pure edit patterns across serial passages (P1-P9). The upper black line and corresponding values indicate the cumulative number of unique pure edit patterns detected from P1 up to each indicated passage, reaching a total of 26,530 unique indels by P9. The area under the curve breaks down the composition at each specific passage: orange represents newly observed pure edit patterns (New Indel) first detected at that time point, while blue represents inherited pure edit patterns (Inherited Indel) shared with prior passages. **(g)** Matrix of pie charts resolving the proportion of newly emerged versus inherited unique pure edit patterns across separate passages and target sites. Rows represent passages P1-P9, and columns indicate individual sgRNA target sites sg1-sg24. Each pie chart details the specific composition of unique pure edit patterns (defined solely by their indel structural configuration without site prefixes) at that coordinate. Orange wedges indicate newly observed edit patterns, whereas blue wedges indicate edit patterns detected in previous passages. **(h)** Detection and editing status of the CERAMIC recorder during cellular reprogramming of E13.5 mouse embryonic fibroblasts (MEFs). The stacked bar chart on the left illustrates the percentage of cells with detected barcodes across three independent single-cell RNA-seq replicates (Rep1-Rep3), classified into cells with edited barcodes (dark blue) and those with unedited detected barcodes only (light blue). The pie chart on the right displays the overall composition pooled across all replicates within the across all transcriptome-qualified cells, distinguishing between cells containing edited barcodes, unedited detected barcodes only, and those with transcriptome only. **(i)** Integrity assessment of target array capture across the three single-cell reprogramming replicates. The stacked bar chart on the left quantifies the proportion of cells recovering different numbers of target array (TA) modules (ranging from 1 to 8, color-coded by a pink-to-dark-purple gradient) within each replicate (Rep1-Rep3). The pie chart summarizes the distribution of recovered TA modules across all three replicates. **(j)** Experimental schematic and western blot analysis showing the tissue-specific expression of Cas9 protein across multiple organs. CERAMIC mice were subjected to doxycycline (Dox) treatment delivered via drinking water at a concentration of 2 mg/mL for 2 weeks to induce systemic recording (top schematic). Representative western blot images (bottom) display Cas9 protein expression (∼150 kDa) across the various harvested tissues indicated above, with GAPDH (∼35 kDa) serving as the internal loading control and PC representing the positive control lane. **(k)** Quantification of allele editing efficiencies across diverse organs in the presence or absence of doxycycline. The scatter dot plot illustrates the percentage of edited alleles (%) across the indicated tissues. Light gray circles represent mice without doxycycline treatment (No Dox), and dark gray circles represent doxycycline-treated mice (Dox). Data points represent individual biological replicates, with error bars indicating mean s.e.m.

**Supplementary Figure 2.**
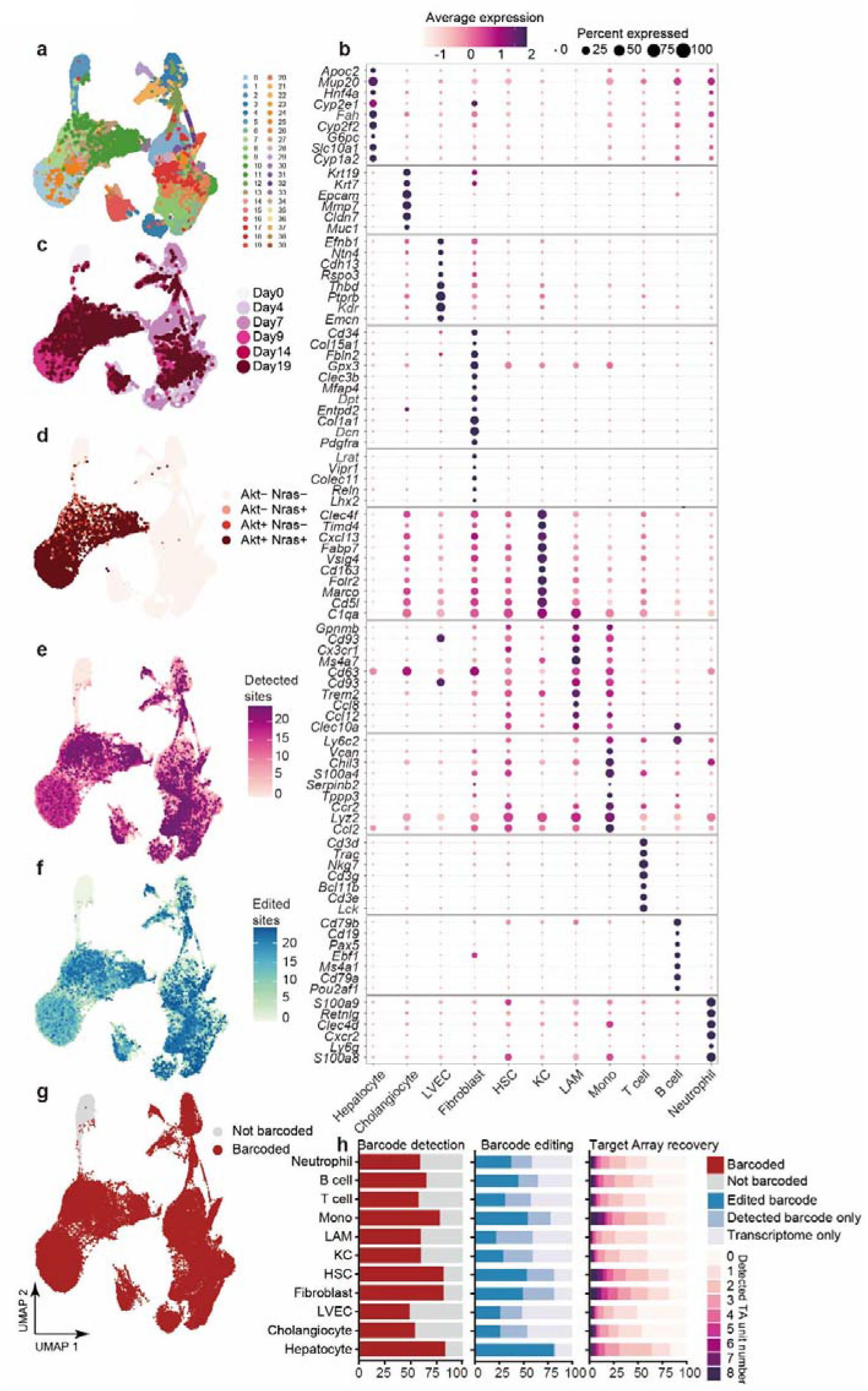
Single-cell atlas construction and CERAMIC barcode recovery in vivo. **(a)** UMAP embedding of the single-cell dataset partitioned into detailed sub-clusters, with color-coding indicating distinct cluster identities (clusters 1 to 38). **(b)** Dot plot showing the expression profiles of canonical cell-type-specific marker genes across the major identified hepatic, stromal, and immune lineages (listed on the horizontal axis). The color intensity of each dot represents the scaled average expression level, and the dot size indicates the percentage of cells within that population expressing the gene. **(c)** UMAP embedding of the cell atlas color-coded by sample collection time points across the progression lineage (Day 0, Day 4, Day 7, Day 9, Day 14, and Day 19), displaying the shifting cellular distribution over time. **(d)** UMAP embedding illustrating the transfection and oncogene integration status of individual cells based on transposon expression. Cells are categorized into four groups: Akt- Nras- (untransfected/wild-type), Akt- Nras+ (N-Ras-only), Akt+ Nras- (Akt-only), and Akt+ Nras+ (double-positive oncogenic hepatocytes). **(e, f)** Spatial quantification of lineage recording performance mapped onto the cell atlas. **(e)** UMAP plot tracking the number of detected target sites per cell, color-coded by a purple-to-pink gradient. **(f)** UMAP plot tracking the number of successfully edited target sites per cell, color-coded by a blue-to-green gradient. **(g)** UMAP embedding colored by lineage-barcode detection status. Cells with detectable barcodes are shown in red, and cells without detectable barcodes are shown in gray. **(h)** CERAMIC barcode detection, barcode editing, and target-array recovery across the indicated cell populations. Left (Barcode detection): Stacked bar chart showing the proportions of barcoded (red) and non-barcoded (gray) cells for each cell type. Middle (Barcode editing): Stacked bar chart showing the proportions of cells containing edited barcodes (dark blue), detected but unedited barcodes (light blue), or transcriptome information only (light purple). Right (Target-array recovery): Stacked bar chart showing the distribution of recovered target-array (TA) modules per cell, with colors indicating the number of recovered TA modules (0-8; pink-to-dark-purple gradient).

**Supplementary Figure 3.**
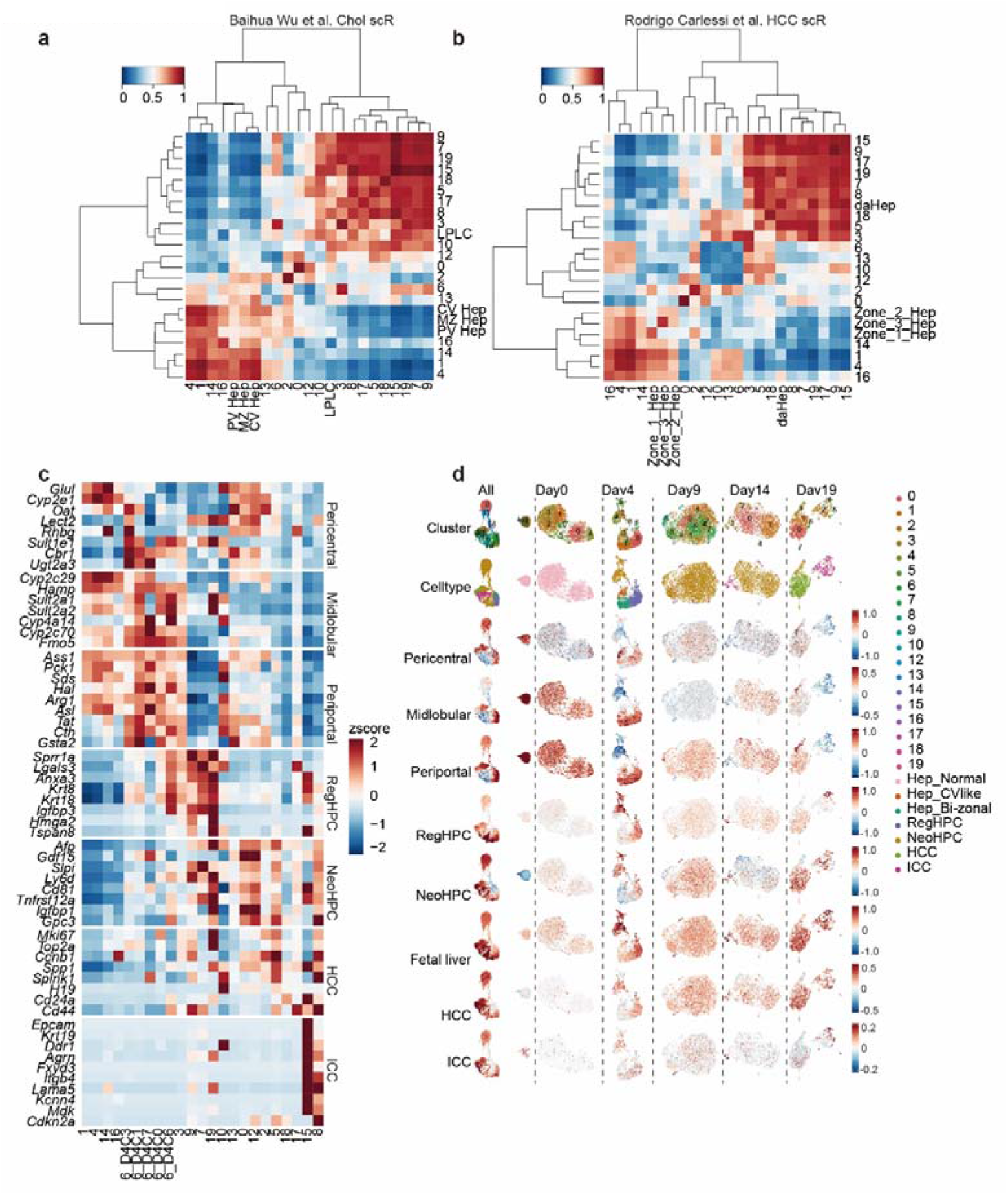
External validation of hepatocyte-state annotations. **(a)** Unsupervised hierarchical clustering and correlation heatmap comparing the hepatocyte sub-clusters identified in this study with hepatocyte populations from a published cholestatic liver injury dataset (Baihua Wu *et al.*). Rows and columns represent hepatocyte sub-clusters identified in this study (clusters 0-19) and reference hepatocyte populations, including periportal (PV Hep), mid-zone (MZ Hep), pericentral (CV Hep) hepatocytes, and liver progenitor-like cells (LPLC). Color indicates Pearson correlation coefficients. **(b)** Unsupervised hierarchical clustering and correlation heatmap comparing the hepatocyte sub-clusters identified in this study with hepatocyte populations from a published fatty liver-associated liver cancer dataset (Rodrigo Carlessi *et al.*). Rows and columns represent hepatocyte sub-clusters identified in this study (clusters 0-19) and reference hepatocyte populations, including Zone_1_Hep, Zone_2_Hep, Zone_3_Hep, and disease-associated hepatocytes (daHep). Color indicates Pearson correlation coefficients. **(c)** Heatmap showing the scaled expression (z-score) of representative marker genes across all identified hepatocyte sub-clusters (columns, clusters 0-19). Genes are grouped according to pericentral, midlobular, periportal, RegHPC, NeoHPC, HCC, and ICC marker sets. Color indicates relative scaled expression. **(d)** UMAP visualization of hepatocyte sub-clusters, cell-state annotations, and signature module scores across the integrated dataset and individual time points. Rows show sub-cluster identities (Cluster; clusters 0-19), annotated hepatocyte states (Celltype), and module scores for pericentral, midlobular, periportal, RegHPC, NeoHPC, fetal liver, HCC, and ICC signatures. Columns show the integrated dataset (“All”) and individual time points (Day 0, Day 4, Day 9, Day 14, and Day 19). Colors indicate relative module scores.

**Supplementary Figure 4.**
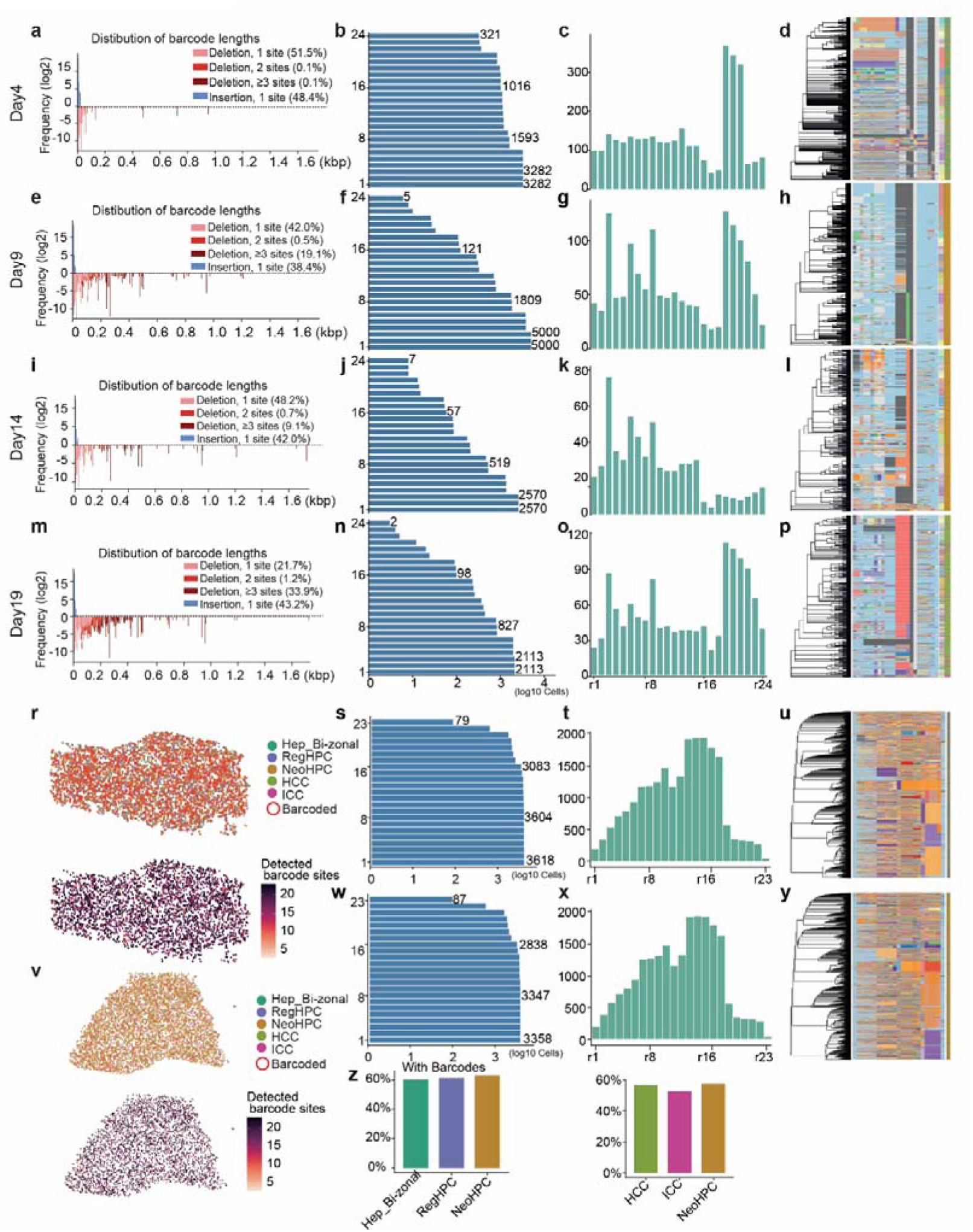
Recovery of lineage information in single-cell and spatial transcriptomic datasets. **(a-d)** Day 4 scRNA-seq dataset. **(a)** Frequency distribution of editing events. The y-axis shows the log₂-transformed frequency of editing events, the x-axis indicates insertion or deletion size (bp), and colors distinguish single-site deletions, multi-site deletions, and single-site insertions. **(b)** Horizontal bar plot showing the cumulative distribution of cells (log₁₀ cells) with at least 1 (≥1) to 24 (≥24) detected target sites; representative cell numbers are indicated. **(c)** Bar chart showing the number of unique editing patterns at individual target sites (r1-r24). **(d)** Single-cell phylogenetic tree with the corresponding character matrix. Rows represent individual cells, colors denote lineage scar states, dark gray indicates missing data, and light blue indicates wild-type alleles (WT). **(e-h)** Day 9 scRNA-seq dataset. **(e)** Frequency distribution of editing events. **(f)** Horizontal bar plot showing the cumulative distribution of cells (log₁₀ cells) with at least 1 (≥1) to 24 (≥24) detected target sites. **(g)** Bar chart showing the number of unique editing patterns at individual target sites (r1-r24). **(h)** Single-cell phylogenetic tree with the corresponding character matrix (dark gray, missing data; light blue, WT). **(i-l)** Day 14 scRNA-seq dataset. **(i)** Frequency distribution of editing events. **(j)** Horizontal bar plot showing the cumulative distribution of cells (log₁₀ cells) with at least 1 (≥1) to 24 (≥24) detected target sites. **(k)** Bar chart showing the number of unique editing patterns at individual target sites (r1-r24). **(l)** Single-cell phylogenetic tree with the corresponding character matrix (dark gray, missing data; light blue, WT). **(m-p)** Day 19 scRNA-seq dataset. **(m)** Frequency distribution of editing events. **(n)** Horizontal bar plot showing the cumulative distribution of cells (log₁₀ cells) with at least 1 (≥1) to 24 (≥24) detected target sites. **(o)** Bar chart showing the number of unique editing patterns at individual target sites (r1-r24). **(p)** Single-cell phylogenetic tree with the corresponding character matrix (dark gray, missing data; light blue, WT). **(r-u)** Day 9 spatial transcriptomic dataset. **(r)** Spatial maps showing tissue coordinates. The upper panel shows spots colored by cell state, with red outlines indicating spots in which lineage barcodes were detected. The lower panel shows the number of detected target sites per spot, colored according to the indicated purple-to-pink scale. **(s)** Horizontal bar plot showing the cumulative distribution of spots (log₁₀) with at least 1 (≥1) to 24 (≥24) detected target sites. **(t)** Bar chart showing the number of unique editing patterns at individual target sites (r1-r24). **(u)** Spatial phylogenetic tree with the corresponding character matrix (dark gray, missing data; light blue, WT). **(v-y)** Day 14 spatial transcriptomic dataset. **(v)** Spatial maps showing tissue coordinates. The upper panel shows spots colored by cell state, with red outlines indicating spots in which lineage barcodes were detected. The lower panel shows the number of detected target sites per spot, colored according to the indicated scale. **(w)** Horizontal bar plot showing the cumulative distribution of spots (log₁₀) with at least 1 (≥1) to 24 (≥24) detected target sites. **(x)** Bar chart showing the number of unique editing patterns at individual target sites (r1-r24). **(y)** Spatial phylogenetic tree with the corresponding character matrix (dark gray, missing data; light blue, WT). **(z)** Bar charts showing the fraction of cells with recovered lineage barcodes across the indicated cell states in the two spatial transcriptomic datasets. The left panel shows Hep_Bi-zonal, RegHPC, and NeoHPC cells, and the right panel shows HCC, ICC, and NeoHPC cells.

**Supplementary Figure 5.**
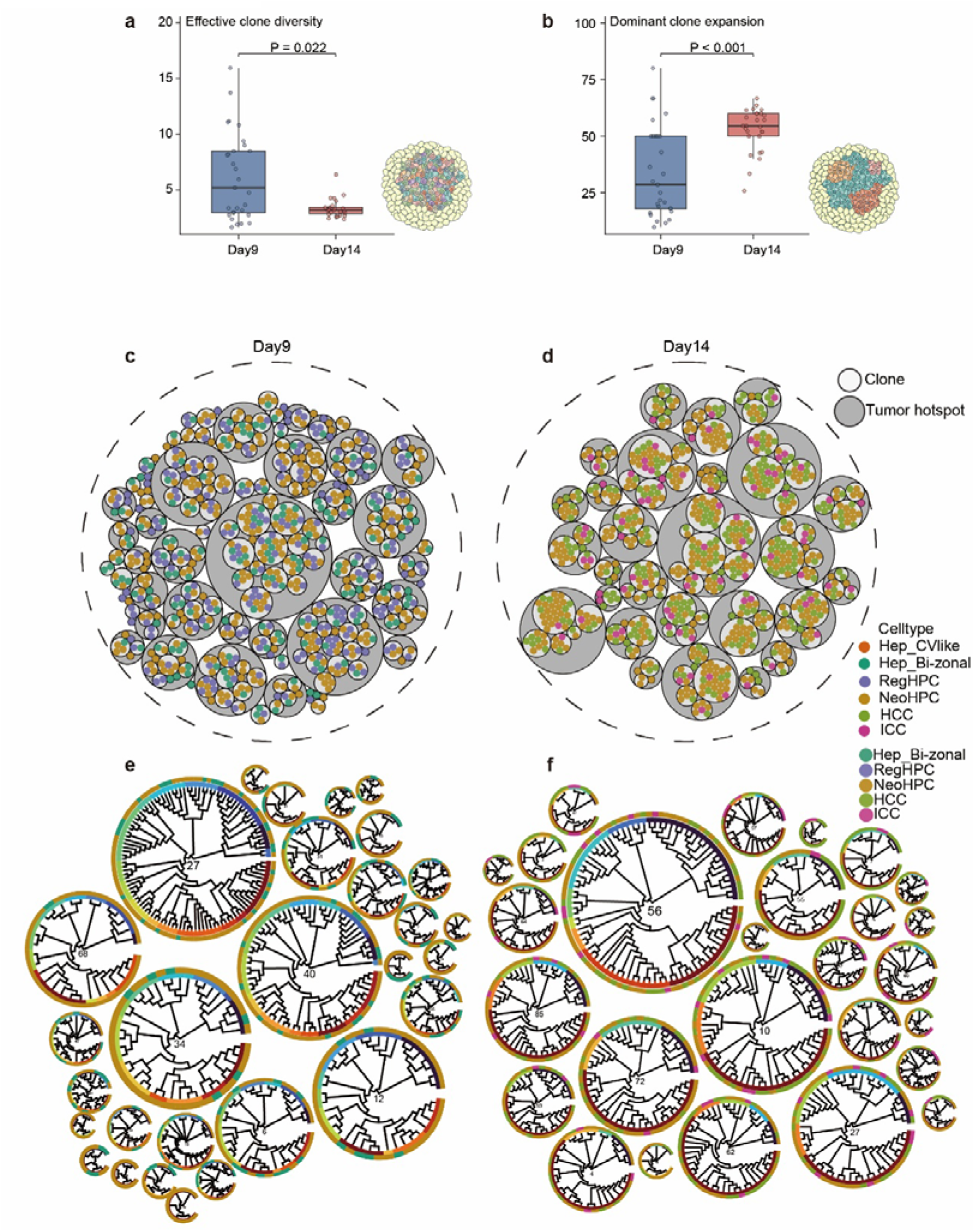
Quantitative clonal dynamics and phylogenetic architecture within spatially restricted tumor hotspots. **(a)** Box plot showing the effective clone number of individual tumor hotspots at Day 9 and Day 14 after induction. Each dot represents one tumor hotspot. The exact *P* value is indicated. The adjacent schematic illustrates the corresponding hotspot organization. **(b)** Box plot showing the fraction of cells contributed by the dominant clone within individual tumor hotspots at Day 9 and Day 14. Each dot represents one tumor hotspot. The exact *P* value is indicated. The adjacent schematic illustrates the corresponding hotspot organization. **(c, d)** Circle-packing visualization of tumor hotspots and reconstructed clones at Day 9 (c) and Day 14 (d). The outer dashed boundary indicates the analyzed tissue region, gray circles represent individual tumor hotspots, white-outlined circles represent reconstructed clones, and each dot represents a single cell colored according to its annotated cell state. **(e, f)** Circular phylogenetic trees of reconstructed clones within individual tumor hotspots at Day 9 (e) and Day 14 (f). Each circular tree corresponds to one tumor hotspot, with inner numbers indicating the hotspot index. The inner ring indicates clone assignments, and the outer ring indicates annotated cell states.

**Supplementary Figure 6.**
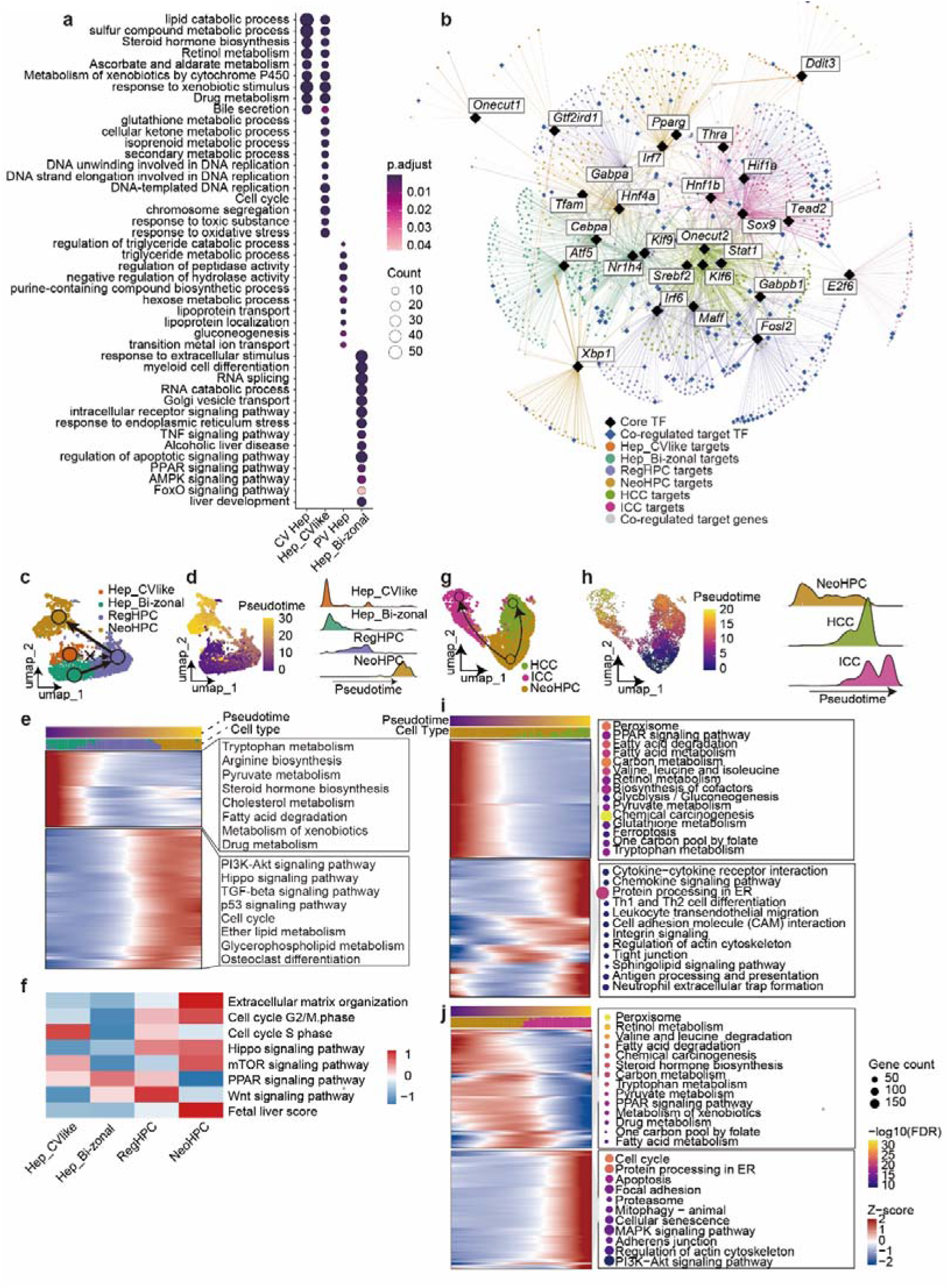
Pseudotime and regulatory analyses of hepatocyte-state transitions. **(a)** Dot plot comparing functional pathway enrichment across normal reference and tumor-fated hepatocyte populations. Marker genes identified in normal pericentral (CV Hep) and periportal (PV Hep) hepatocytes from the reference dataset (Baihua Wu et al.) are aligned against the marker genes of Hep_CVlike and Hep_Bi-zonal states from the current dataset to evaluate functional conservation and divergence during early tumor initiation. Dot size indicates the gene count contributing to each term, and dot color indicates the adjusted significance (P-adjust). **(b)** Hub transcription factor-centered regulatory network highlighting convergent transcriptional programs. This plot serves as an expanded, highly detailed visualization of the core regulatory wiring shown in Figure 4d. Black diamonds denote master hub TFs, blue diamonds represent downstream co-regulated target genes that function as independent TFs, and small circles indicate non-TF target genes. Edges and target nodes are color-coded based on their cell-state assignments (Hep_CVlike, Hep_Bi-zonal, RegHPC, NeoHPC, HCC, and ICC). Targets shared by multiple master regulators are prioritized toward the center to highlight coordinated downstream transcriptional programs, while high-weight private targets are positioned at the periphery to preserve TF-specific local structures. **(c)** UMAP embedding overlaying the cellular lineage relationships and differentiation paths derived from lineage-barcode scar recordings among early pre-malignant populations (Hep_CVlike, Hep_Bi-zonal, RegHPC, and NeoHPC). **(d)** Pseudotime trajectory of early pre-malignant states. Left panel: UMAP plot color-coded by the continuous pseudotemporal gradient inferred via trajectory analysis. Right panel: Ridge plot displaying the continuous cell density distribution of each cell state along the progression pseudotime axis. **(e)** Gene expression dynamics and functional rewiring along the pre-malignant pseudotime trajectory. The heatmap illustrates the continuous cascade of transition-associated genes ordered by their peak expression along pseudotime (top horizontal bar). Aligned right panels display representative enriched KEGG pathways corresponding to distinct temporal gene blocks. **(f)** Heatmap displaying pathway module scores for liver injury-associated signaling cascades and the curated Fetal Liver Score compared across the four pre-malignant cell states (Hep_CVlike, Hep_Bi-zonal, RegHPC, and NeoHPC). The relative score values are standardized and clipped (z-score color scale). **(g)** UMAP embedding mapping the terminal lineage relationships and branching architecture determined by lineage barcode evidence during advanced malignant progression from NeoHPC precursors into divergent HCC and ICC terminal fates. **(h)** Pseudotime trajectory of terminal malignant bifurcation. Left panel: UMAP visualization displaying the pseudotime trajectory originating from NeoHPC cells. Right panel: Ridge plot illustrating the distinct cell density distributions of NeoHPC, HCC, and ICC populations across the terminal pseudotime spectrum. **(i, j)** Bifurcated gene expression cascades and pathway profiles tracking the divergence from NeoHPC toward HCC **(i)** and ICC **(j)** fates. The heatmaps display scaled gene expression kinetics along their respective lineage-specific pseudotime paths. Adjacent dot plots summarize the corresponding KEGG pathway enrichment terms for each sequential gene module, with dot size representing the gene count and dot color indicating significance (-log FDR).

**Supplementary Figure 7.**
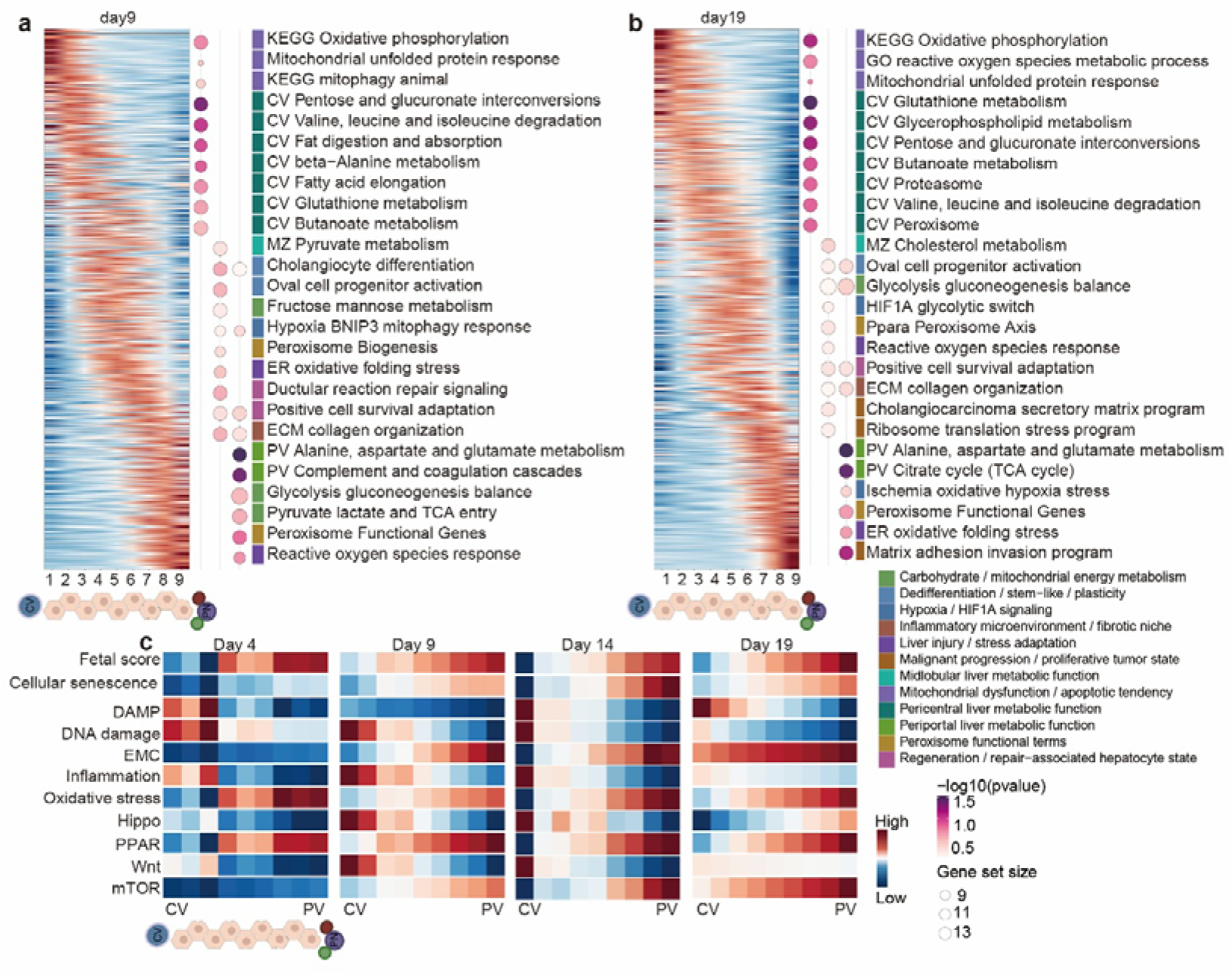
Spatial pathway remodeling along the liver zonation axis. **(a,b)** Spatial pathway analysis across nine concentric layers in tumor regions at Day 9 (a) and Day 19 (b). Spatial transcriptomic cells were grouped into nine layers according to their relative position along the lobular axis, with layers 1-3 corresponding to the pericentral (CV) region, layers 4-6 to the midlobular region, and layers 7-9 to the periportal (PV) region. Left, heatmaps showing scaled expression of genes ordered according to their spatial layer preference. Rows represent selected genes, columns represent the mean expression of each spatial layer, and colors indicate relative expression (red, higher; blue, lower). Right, dot plots showing representative enriched GO and KEGG pathways for the pericentral, midlobular, and periportal layer groups. Pathways are color-coded according to the functional categories shown in Figure 4a. Dot size indicates the number of genes in each pathway, and dot color indicates enrichment significance (-log₁₀ *P* value). **(c)** Heatmap showing pathway module scores across nine spatial layers from the pericentral (CV, layer 1) to the periportal (PV, layer 9) region at Day 4, Day 9, Day 14, and Day 19. Rows represent pathway modules, including fetal liver, cellular senescence, DAMP, DNA damage, Hippo, and Wnt signatures. Columns represent the indicated spatial layers at each time point. Values are displayed as standardized z-scores, with red indicating relatively higher pathway scores and blue indicating relatively lower pathway scores.

**Supplementary Figure 8.**
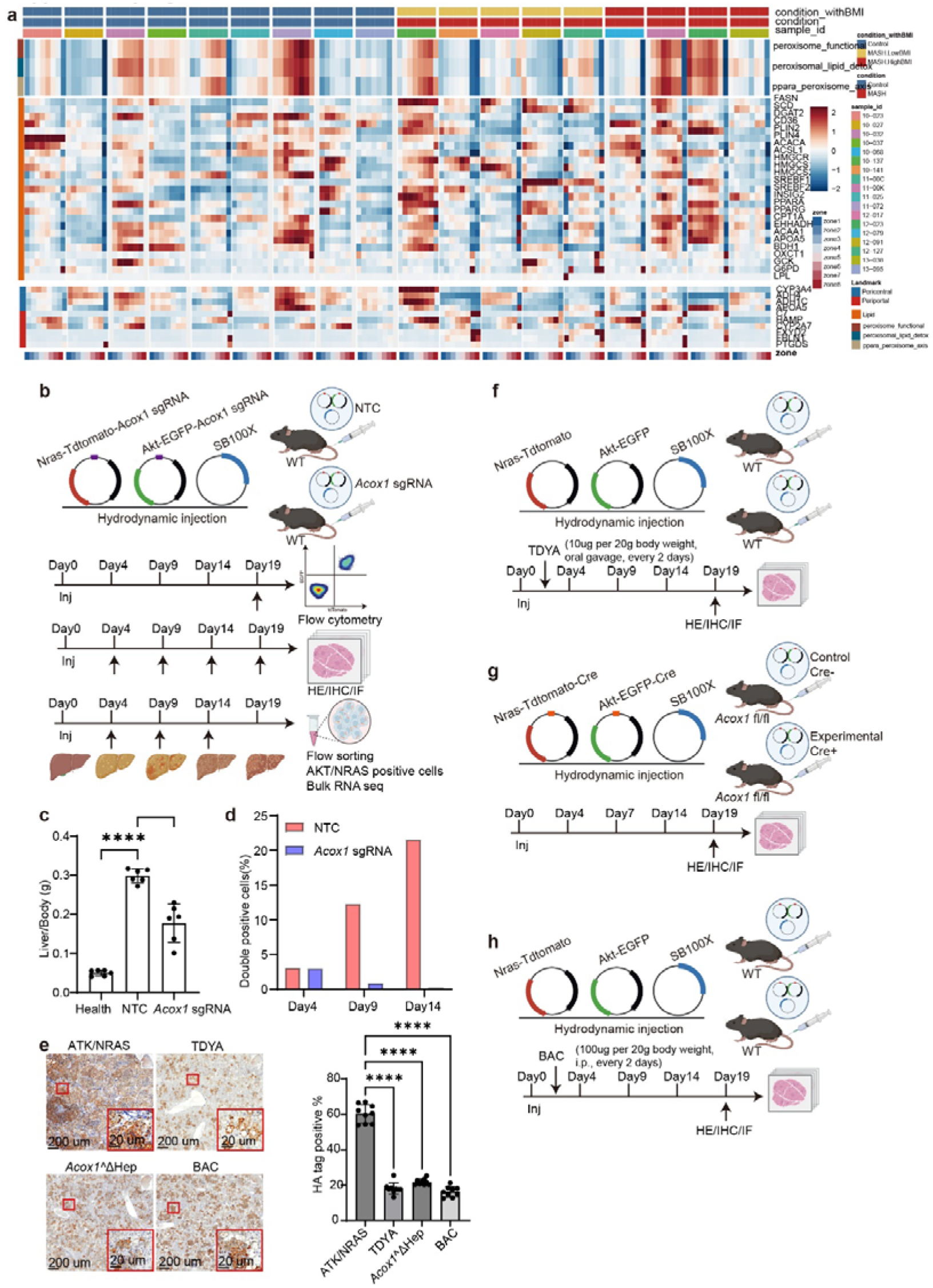
Experimental design and validation of Acox1 perturbation. **(a)** Heatmap showing peroxisomal and metabolic gene and pathway module scores in the Xenium spatial transcriptomic dataset shown in Figure 5b. Columns are grouped by individual patient, with eight columns representing spatial zones 1-8 for each patient. The top annotation bars indicate patient group and BMI category (blue, Control; yellow, MASH-lowBMI; red, MASH-highBMI), disease condition (blue, Control; red, MASH), and patient identifier. Rows represent genes or pathway modules. The upper three rows correspond to peroxisome-related gene sets, the middle block marked by the orange side annotation contains lipid metabolism-associated genes, and the lower rows contain pericentral-associated and periportal-associated metabolic zonation markers, indicated by blue and red side annotations, respectively. **(b)** Experimental timeline and vector design for CRISPR-mediated *Acox1* perturbation in the AKT/NRAS model. Vector diagrams show Nras-tdTomato-*Acox1* sgRNA (tdTomato, red; *Acox1* sgRNA, purple; Nras, black), Akt-EGFP-*Acox1* sgRNA (EGFP, green; Akt, black), and the SB100X transposase vector (transposase, blue). Samples were collected at the indicated time points for flow cytometry, histological analyses, including H&E, immunohistochemistry, and immunofluorescence, and fluorescence-activated cell sorting of oncogene-expressing cells for bulk RNA-seq. **(c)** Quantification of the liver-to-body-weight ratio corresponding to the representative gross liver images shown in Figure 5d. Data were analyzed by one-way ANOVA followed by Tukey’s multiple-comparisons test (n = 6 mice per group; healthy versus NTC, adjusted *P* < 0.0001; healthy versus *Acox1* sgRNA, adjusted *P* < 0.0001; NTC versus *Acox1* sgRNA, adjusted *P* < 0.0001). **(d)** Percentage of EGFP⁺tdTomato⁺ cells at Days 4, 9, and 14 after induction. Red denotes the non-targeting control (NTC) group, and blue denotes the *Acox1* sgRNA group. The indicated gates were used for fluorescence-activated cell sorting of EGFP⁺tdTomato⁺ cells for the bulk RNA-seq analysis shown in Figure 5h. **(e)** Representative HA immunohistochemical images and quantification for untreated AKT/NRAS, TDYA-treated, hepatocyte-specific *Acox1*-deleted (*Acox1*^ΔHep^), and BAC-treated groups. HA immunostaining detects HA-tagged AKT-expressing cells. Scale bars, 200 μm; enlarged views, 20 μm. Data were analyzed by one-way ANOVA with Dunnett’s multiple-comparisons test (n = 9 fields of view per group; AKT/NRAS versus TDYA, adjusted *P* < 0.0001; AKT/NRAS versus *Acox1*^ΔHep^, adjusted *P* < 0.0001; AKT/NRAS versus BAC, adjusted *P* < 0.0001). **(f)** Experimental timeline and vector design for pharmacological inhibition of ACOX1 with TDYA in AKT/NRAS-induced mice. TDYA was administered by oral gavage at 10 μg per 20 g body weight every other day. **(g)** Experimental strategy for hepatocyte-specific deletion of *Acox1* using *Acox1*^fl/fl^ mice and Cre-expressing oncogenic vectors, including Nras-tdTomato-Cre, Akt-EGFP-Cre, and SB100X. **(h)** Experimental timeline for pharmacological inhibition of P4HB with bacitracin (BAC) in AKT/NRAS-induced mice. BAC was administered by intraperitoneal injection at 100 μg per 20 g body weight every other day.

**Supplementary Figure 9.**
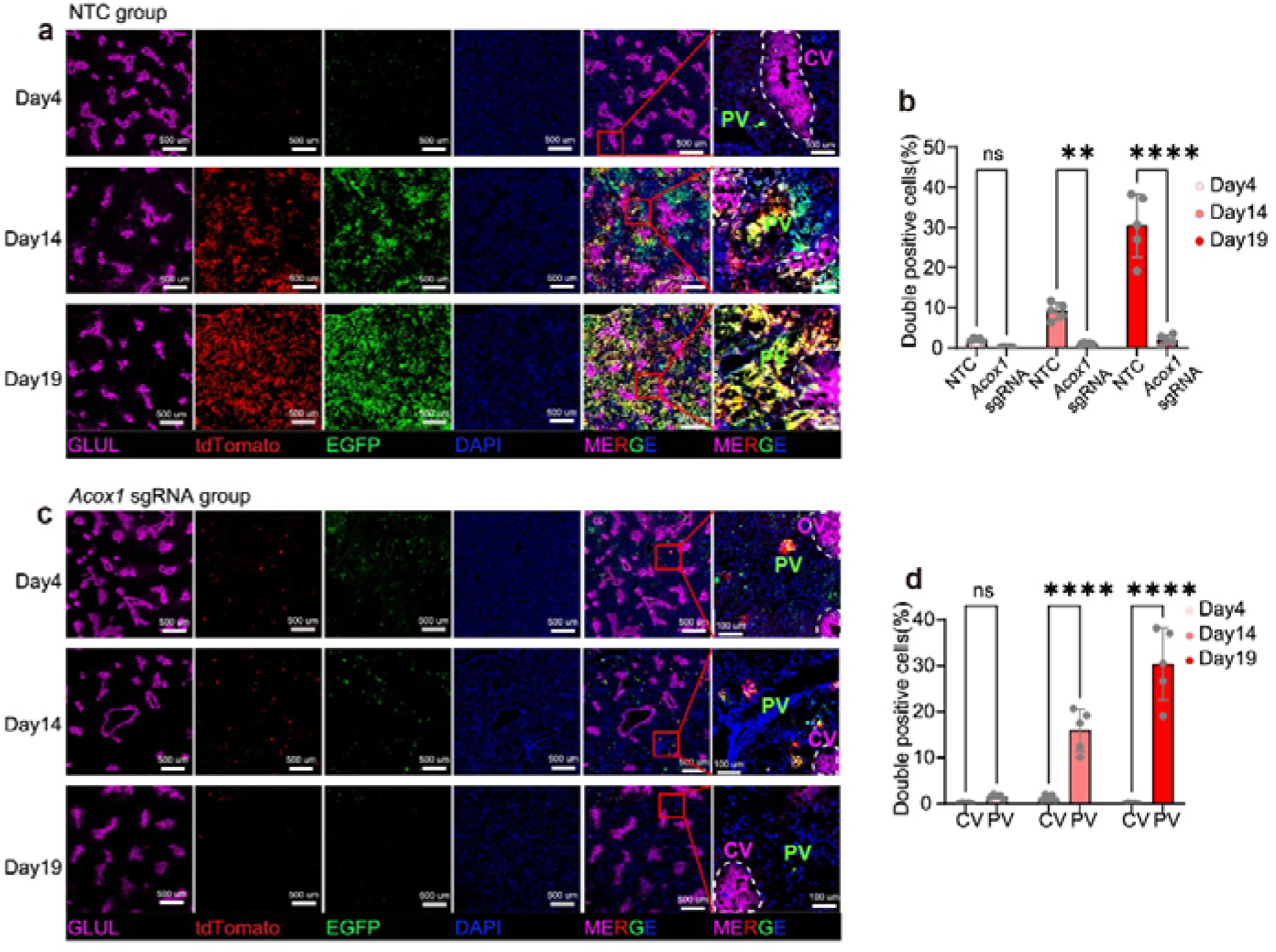
Acox1 perturbation restricts transformed-cell expansion outside the pericentral region. **(a)** Representative multiplex immunofluorescence images of liver sections from non-targeting control (NTC) mice at Days 4, 14, and 19 after induction. Separate and merged channels are shown for GLUL (magenta), tdTomato (red), EGFP (green), and DAPI (blue). Enlarged views of the pericentral (CV, outlined by white dashed lines) and periportal (PV) regions are shown on the right. Scale bars, 500 μm (main images) and 100 μm (enlarged views). **(b)** Quantification of EGFP⁺tdTomato⁺ cells in NTC and *Acox1* sgRNA-treated mice at Days 4, 14, and 19. Data were analyzed by two-way ANOVA with Sidak’s multiple-comparisons test (n = 6 fields of view per group per time point; Day 4, adjusted *P* = 0.7870; Day 14, adjusted *P* < 0.0001; Day 19, adjusted *P* < 0.0001). **(c)** Representative multiplex immunofluorescence images of liver sections from *Acox1* sgRNA-treated mice at Days 4, 14, and 19 after induction. Separate and merged channels are shown for GLUL (magenta), tdTomato (red), EGFP (green), and DAPI (blue). Enlarged views of the pericentral (CV, outlined by white dashed lines) and periportal (PV) regions are shown on the right. Scale bars, 500 μm (main images) and 100 μm (enlarged views). **(d)** Quantification of EGFP⁺tdTomato⁺ cells in the pericentral (CV) and periportal (PV) regions at Days 4, 14, and 19. Data were analyzed by two-way ANOVA with Sidak’s multiple-comparisons test (n = 6 fields of view per region per time point; Day 4, adjusted *P* = 0.9138; Day 14, adjusted *P* < 0.0001; Day 19, adjusted *P* < 0.0001).

**Supplementary Figure 10.**
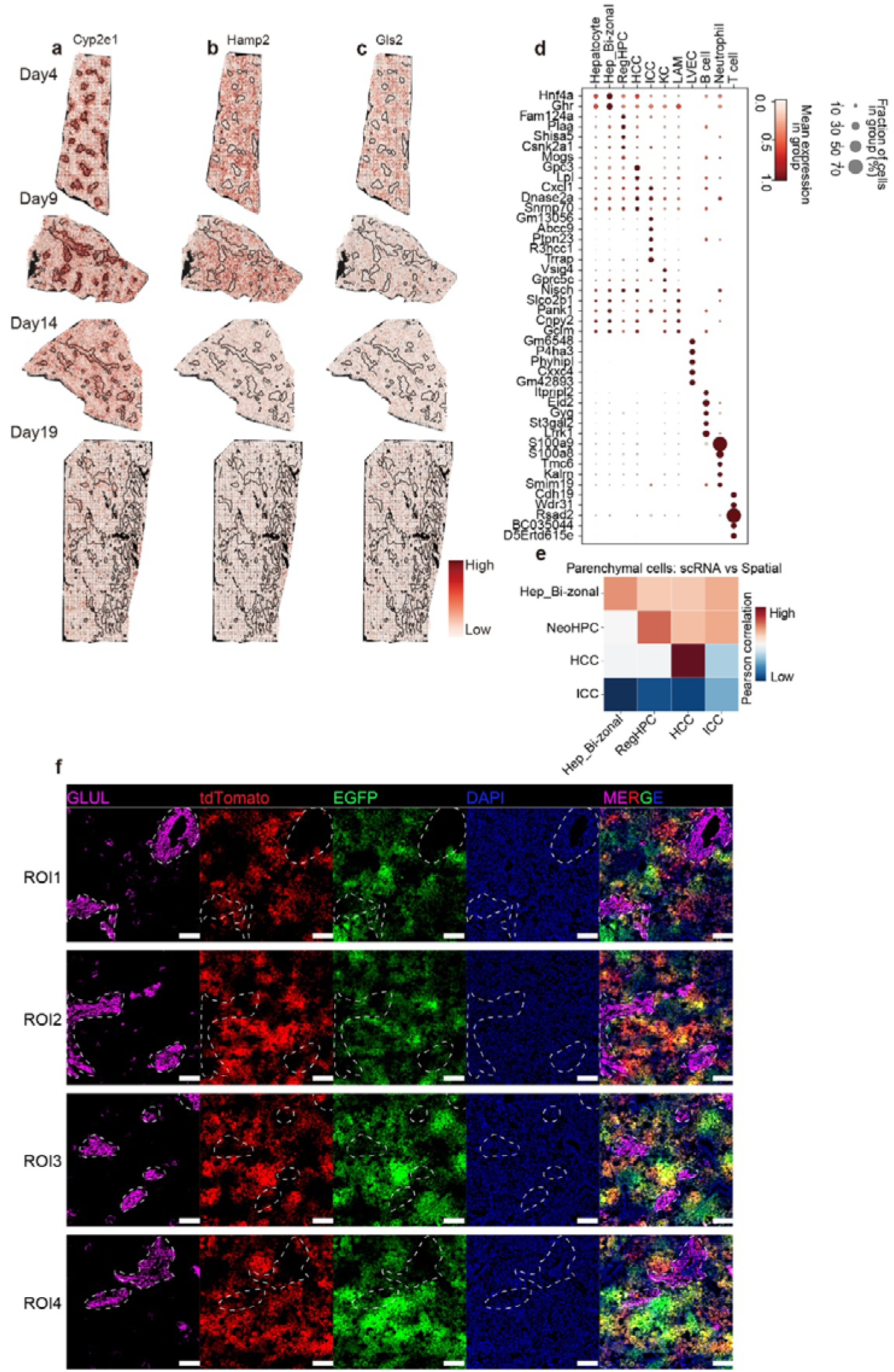
Spatial validation of zonation and cell-state annotations. **(a-c)** Spatial expression maps of representative liver zonation markers at Days 4, 9, 14, and 19 after induction. Spatial plots show the expression of **(a)** *Cyp2e1*, **(b)** *Hamp2*, and **(c)** *Gls2*, representing pericentral-, midlobular-, and periportal-associated markers, respectively. Black outlines indicate the pericentral regions. Colors indicate relative expression levels from low (light pink) to high (dark red). **(c)** Dot plot showing representative marker genes used to annotate parenchymal, stromal, and immune cell populations in the spatial transcriptomic dataset. Rows represent the indicated cell populations, including hepatocytes, Hep_Bi-zonal, RegHPC, HCC, ICC, Kupffer cells (KC), lipid-associated macrophages (LAM), endothelial cells (LVEC), B cells, neutrophils, and T cells. Dot size indicates the percentage of cells expressing each gene, and dot color indicates the relative mean expression level. **(d)** Heatmap showing Pearson correlation coefficients between cell-state expression profiles derived from single-cell RNA sequencing (scRNA-seq; rows) and spatial transcriptomics (columns). The indicated cell states include Hep_Bi-zonal, NeoHPC, HCC, and ICC. Colors indicate Pearson correlation coefficients from low (blue) to high (red). **(e)** Representative multiplex immunofluorescence images from four independent regions of interest (ROI1-ROI4) at Day 19 after induction. Separate and merged channels are shown for GLUL (magenta), tdTomato (red), EGFP (green), and DAPI (blue). White dashed lines indicate GLUL-defined pericentral (CV) regions. Scale bars, 200 μm.

**Supplementary Figure 11.**
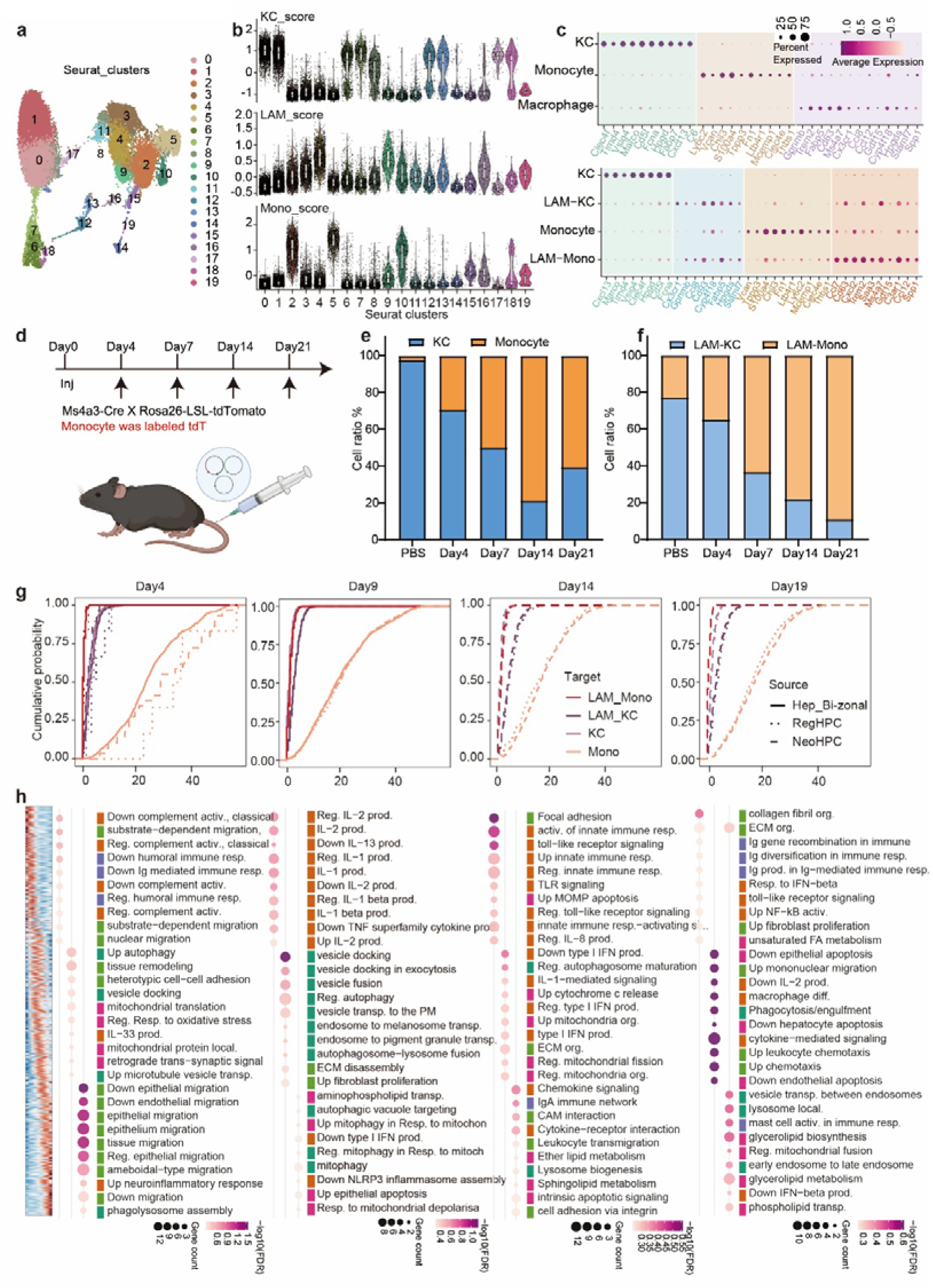
Myeloid annotation, monocyte lineage tracing and spatial macrophage interactions. **(a)** UMAP visualization of hepatic myeloid cell populations showing 20 transcriptionally defined subclusters. **(b)** Violin plots showing signature scores for Kupffer cells (KC, top), lipid-associated macrophages (LAM, middle), and monocytes (bottom) across the indicated myeloid subclusters. **(c)** Dot plots showing representative marker genes across major myeloid populations (top, KC, monocytes, and macrophages) and lipid-associated macrophage populations (bottom, KC, LAM-KC, monocytes, and LAM-Mono). Dot size indicates the percentage of cells expressing each gene, and dot color indicates the relative mean expression level. **(d)** Experimental strategy using *Ms4a3*-Cre; *Rosa26*-LSL-tdTomato mice combined with hydrodynamic injection of oncogenic vectors to label bone marrow-derived monocytes during liver tumor initiation. **(e, f)** Quantification of the proportions of myeloid cell populations at baseline (PBS) and Days 4, 7, 14, and 21 after induction. **(e)** Kupffer cells (KC) and monocytes. **(f)** LAM-KC and LAM-Mono populations. Data were analyzed using two-way repeated-measures ANOVA for (e) (interaction, *P* < 0.0001; column factor, *P* < 0.0001) and two-way repeated-measures ANOVA with Geisser-Greenhouse correction for (f) (interaction, *P* < 0.0001; column factor, *P* < 0.0001). **(f)** Nearest-neighbor distance analysis between hepatocyte states and myeloid cell populations at Days 4, 9, 14, and 19. Curves show the cumulative probability distribution as a function of the distance between Hep_Bi-zonal (solid line), RegHPC (dotted line), or NeoHPC (dashed line) cells and the indicated myeloid populations (LAM-Mono, LAM-KC, KC, and monocytes). **(g)** Spatial analysis of lipid-associated macrophages (LAMs) across lobular layers. Spatial transcriptomic LAMs were grouped according to their relative position along the lobular axis from the pericentral (CV-like) to the periportal (PV-like) region. Left, heatmap showing scaled expression of genes ordered according to their spatial layer preference at Day 4. Rows represent selected genes, columns represent the mean expression of each spatial layer, and colors indicate relative expression levels. Right, dot plots showing representative enriched GO and KEGG pathways across Days 4, 9, 14, and 19 based on spatial layer-associated genes. Pathways are grouped into the same functional categories as those shown in Figure 7e, including inflammation/innate immune response, antigen presentation/immune regulation, phagocytosis/endocytosis, and metabolism/lipid-related categories. Dot size indicates the number of genes associated with each pathway, and dot color indicates enrichment significance (-log₁₀ FDR).

